# Using neural biomarkers to personalize dosing of vagus nerve stimulation

**DOI:** 10.1101/2023.08.30.555487

**Authors:** Antonin Berthon, Lorenz Wernisch, Myrta Stoukidi, Michael Thornton, Olivier Tessier-Lariviere, Pascal Fortier-Poisson, Jorin Mamen, Max Pinkney, Susannah Lee, Elvijs Sarkans, Luca Annecchino, Ben Appleton, Philip Garsed, Bret Patterson, Samuel Gonshaw, Matjaz Jakopec, Sudhakaran Shunmugam, Tristan Edwards, Aleksi Tukiainen, Joel Jennings, Guillaume Lajoie, Emil Hewage, Oliver Armitage

## Abstract

**Background:** Vagus nerve stimulation (VNS) is an established therapy for treating a variety of chronic diseases, such as epilepsy, depression, obesity, and for stroke rehabilitation. However, lack of precision and side-effects have hindered its efficacy and extension to new conditions.

**Objective:** To achieve a better understanding of the relationship between VNS parameters and neural and physiological responses to enable the design of personalized dosing procedures to improve precision and efficacy of VNS therapies.

**Methods:** We used biomarkers from recorded evoked neural activity and short-term physiological responses (throat muscle, cardiac and respiratory activity) to understand the response to a wide range of VNS parameters in anaesthetised pigs. Using signal processing, Gaussian processes (GP) and parametric regression models we analyse the relationship between VNS parameters and neural and physiological responses.

**Results:** Firstly, we observe inter-subject variability for both neural and physiological responses. Secondly, we illustrate how considering multiple stimulation parameters in VNS dosing can improve the efficacy and precision of VNS therapies. Thirdly, we describe the relationship between different VNS parameters and the evoked neural activity and show how spatially selective electrodes can be used to improve fibre recruitment. Fourthly, we provide a detailed exploration of the relationship between the activations of neural fibre types and different physiological effects, and show that recordings of evoked neural activity are powerful biomarkers for predicting the short-term physiological effects of VNS. Finally, based on these results, we discuss how recordings of evoked neural activity can help design VNS dosing procedures that optimize short-term physiological effects safely and efficiently.

**Conclusion:** Understanding of evoked neural activity during VNS provide powerful biomarkers that could improve the precision, safety and efficacy of VNS therapies.

## 1 Introduction

### 1.1 Therapeutic potential and biological problem

The vagus nerve innervates most visceral organs in the thorax and abdomen, including the pharynx, larynx, heart, lungs, and gut (Johnson and Wilson, 2018), as well as areas within the brain and central nervous system (Thompson, Mastitskaya, and Holder, 2019). As such, it modulates functions such as respiration, circulation, and digestion. Abnormal vagus nerve activity has been associated with a variety of non-communicable diseases, including hypertension (Grassi, 2009), heart failure (Kishi, 2012), epilepsy (Ronkainen et al., 2006), diabetes (Liao et al., 1995), cancer (Gidron, De Couck, and De Greve, 2014), inflammatory diseases (Koopman et al., 2016), obesity (Kral, Paez, and Wolfe, 2009) and eating disorders (Loper et al., 2021).

The vagus nerve’s diverse regulatory functions make it a prime target for therapies, most notably vagus nerve stimulation (VNS). VNS induces therapeutic effects by electrically stimulating the vagus nerve. By adjusting stimulation parameters, VNS can activate specific nerve fibres, resulting in different physiological effects. For example, stimulating large, myelinated A-fibres has an anti-epileptic effect (Bao, Zhou, and Luan, 2011); smaller, myelinated B- and possibly unmyelinated C-fibres are related to anti-inflammatory action (Sundman and Olofsson, 2014), and B-fibres influence cardiac function (Qing et al., 2018).

VNS is FDA-approved to treat epilepsy, depression, obesity, and for stroke rehabilitation (U.S. Food and Drug Administration, PMAs P970003, P130019, P210007). It is being investigated as a therapy for heart failure (Zile et al., 2020), hypertension (Ntiloudi et al., 2019), inflammatory conditions (Pavlov and Tracey, 2012; Kessler et al., 2012), traumatic brain injury (TBI) (Bansal et al., 2012), lung injury (Santos et al., 2011; Reys et al., 2013) Alzheimer’s disease (Merrill, Bikson, and Jefferys, 2005), anxiety (George et al., 2008), chronic pain (Chakravarthy et al., 2015), tinnitus (Tyler et al., 2017), rheumatoid arthritis (Koopman et al., 2016), diabetes (Meyers et al., 2016) and obesity (Val-Laillet et al., 2010; Ikramuddin et al., 2014).

While VNS has helped tens of thousands of patients with chronic disease (primarily refractory epilepsy), a persistent challenge is the therapy’s lack of precision. Response rates range between 50-70% (Englot et al., 2015; Toffa et al., 2020; Batson et al., 2022) and side effects are common, including cough, voice alteration, laryngeal spasms, and local pain Ben-Menachem, 2001. Additionally, it can take up to 12 months of adjusting stimulation parameters before peak effectiveness is reached (LivaNova, 2020).

This lack of precision is largely due to the complex morphology of the vagus nerve (Fig. 2) and the fact that fibres associated with side effects tend to respond to stimulation more readily than those associated with desirable therapeutic effects (Fitchett, Mastitskaya, and Aristovich, 2021). While fascicles are organized at least partially by the organs they innervate and the functions they mediate (Jayaprakash et al., 2022), the vagus nerve exhibits extensive branching and merging. Additionally, nerve morphology is unique for each patient and VNS parameter settings need to be adjusted accordingly. In regard to fibre engagement, the large myelinated fibres that innervate the mucosa and muscles of the neck, larynx and pharynx respond much more readily to stimulation (Gold et al., 2016) than most other fibre types (Yoo et al., 2013). This makes it difficult to avoid side effects such as cough, throat pain, dyspnea, and voice alteration (De Ferrari et al., 2017) when inducing a therapeutic response.

### 1.2 Dosing nerve stimulation

When delivering any neurostimulation based therapy, the user must select the right stimulation parameters that achieve the desired therapeutic effects and avoid side effects. This is commonly referred to as dosing the VNS therapy.

Dosing VNS and other neurostimulation therapies is difficult due to the complex landscape of possible stimulation parameters that can be adjusted presenting a ‘curse of dimensionality’ challenge, further exacerbated by inter-patient variability. As the number of parameters increase to achieve a more precise stimulation, the search space of possible parameter sets expands significantly. While optimal settings should be determined for each patient, brute force methods take too long to do safely or practically in the operating room or clinic, and would expose the patient to potentially dangerous side effects.

In order to address the lack of precision of current VNS approaches, many authors have proposed and developed selective VNS (sVNS) approaches including, selectively blocking undesirable nerve signal propagation (anodal block, kilohertz electrical stimulation block) and using novel electrode arrays and pulse shapes (Fitchett, Mastitskaya, and Aristovich, 2021). While sVNS strategies have shown promise preclinically and in limited clinical use (Plachta et al., 2014; Aristovich et al., 2020; Dali et al., 2018; Pěclin et al., 2009; Ojeda et al., 2016) there are technological and scientific hurdles to overcome before these techniques are widely available in clinics. Additionally, it is specifically notable that in all sVNS approaches the stimulation parameter space is extended in order to describe selective stimulations, for example, by defining sets of complex pulse shapes or multiple electrode locations. This further extends the complex dosing problem rather than solving it.

Faced with these problems, existing dosing methods greatly reduce the complexity of the search space for users, but in doing so remove the possibility to find optimal or personalised doses per patient. During dosing sessions, clinicians choose from preset programs or adjust only one parameter—typically current—until there are noticeable side effects such as voice alteration (LivaNova, 2020). As patients habituate to the stimulation, current can be increased. Even within this highly restricted parameter space, it can take 12 months for patients to reach effective stimulation settings.

### 1.3 Objectives and outline of the present study

In this study, we investigate the potential drawbacks of current clinical dosing strategies and assess whether a broader consideration of dosing may resolve them. Such a broader approach takes advantage of the entire stimulation parameter space in order to find VNS parameters inducing a desired physiological response with minimal side effects. To this end, we examine the relationship between VNS parameters, the evoked compound action potentials (eCAPs), and physiological effects (Fig. 1A), with the aim to support the development of new methods to arrive at a personalized dose efficiently. In pursuit of this, we explored a variety of systematic and extensive sets of VNS parameters for their neural and short-term physiological effects on anaesthetised swine (Fig. 1B), the preferred animal model for the human vagus (Settell et al., 2020). In terms of physiological effects, throughout this study we consider dosing in the context of VNS for heart failure. As such, cardiac responses to VNS such as tachycardia and bradycardia are considered ”ON-target effects”, while common side effects of cervical VNS like bradypnea or laryngeal muscle contractions are treated as ”OFF-target effects” that should be mitigated. The method and approaches presented in this work could equally be applied to other vagally mediated autonomic functions with similar OFF-target effects to mitigate. We focus on short-term effects in the range of minutes, as it has been shown that dosing of VNS parameters for this time frame, with consequent stimulations delivered periodically over time, has the potential for improving clinically relevant outcomes (Dusi and De Ferrari, 2021). This is also the time frame over which off-target effects of VNS occur.

**Figure 1:**
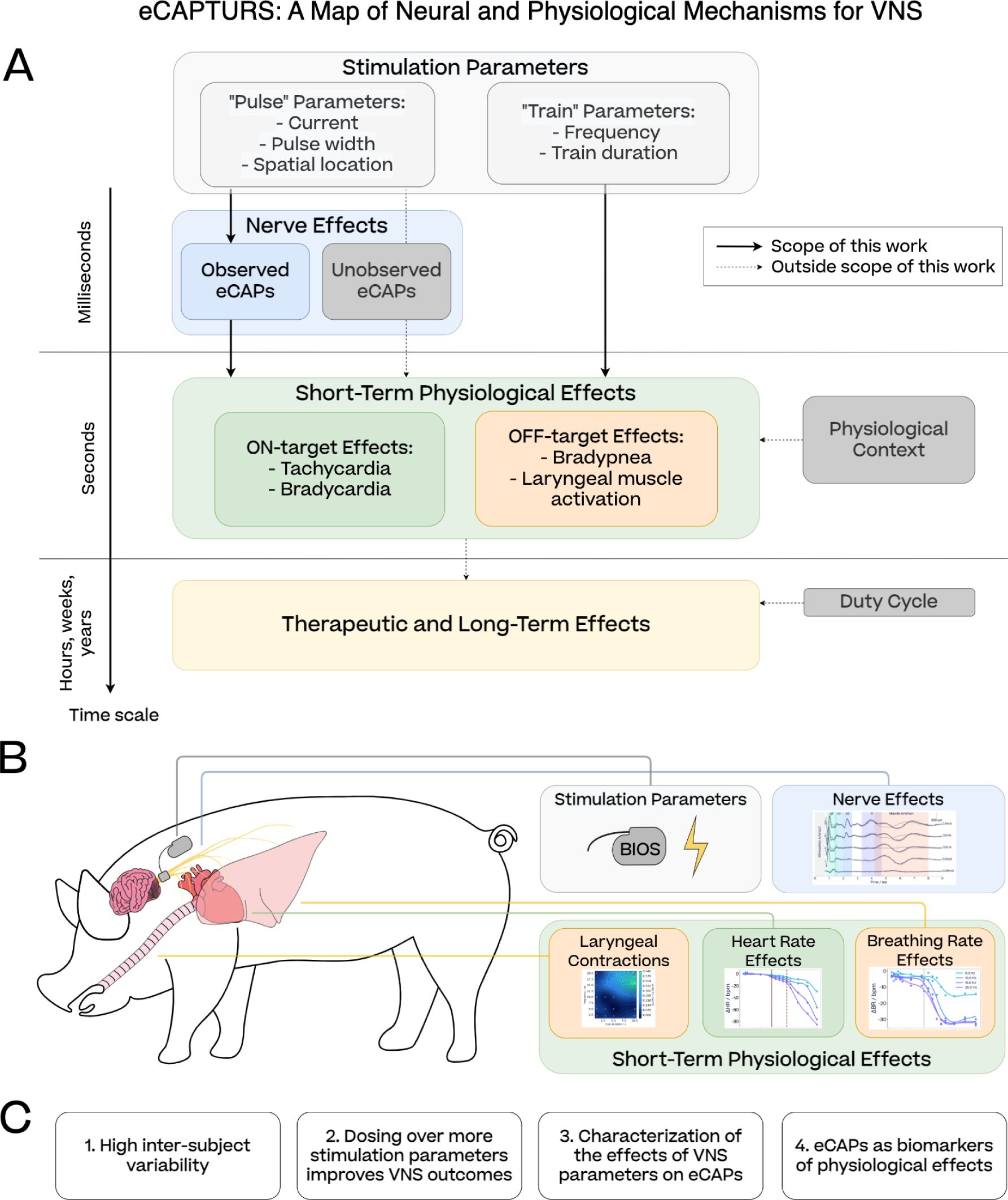
(A) Overview of the eCAPTURS framework (eCAPs To Unravel Responses to Stimulation), showing the relationship between stimulation parameters and responses on the nerve and the physiological level investigated in this study. The three solid arrows represent the main focus of this study. Light grey boxes indicate VNS parameters, as defined in North et al., 2021. The physiological state captures any bodily variables that might affect the short-term responses to stimulations, such as baseline heart rate or breathing rate, timing of the stimulation with respect to the cardiac (Ojeda et al., 2016) or breathing cycle (Sclocco et al., 2019) or anaesthesia (see Section 4.3.4). (B) High-level representation of our VNS setting on porcine subjects: our custom neural interface delivers stimulations to the cervical vagus nerve while evoked neural activity and short-term physiological effects (laryngeal contractions, heart rate and breathing rate changes) are recorded. (C) Key results reported in this study.

Our results fit into four major themes (Fig. 1C). First, we found high inter-subject variability of physiological responses to VNS, which is consistent with observations in current clinical applications of VNS and emphasizes the need for personalized VNS dosing. Second, we find that a search across the entire stimulation space is more successful in producing desirable effects and avoiding side-effects than dosing protocols limited to one or a few parameters. Third, only a subset of VNS parameters, which we term *pulse parameters*, determine the profile of neural fibres activated by a stimulation (Fig. 1A). Another subset, which we term *train parameters*, are responsible for integrating eCAPs into physiological effects. Fourth, we investigate how the causal relationship of eCAPs to physiological effects, as well as their rapid emergence following stimulations, makes eCAPs a highly promising tool for VNS dosing as biomarkers of physiological effects.

These key results suggest two clear applications which we will discuss in Section 4–the first practical, the second theoretical. The first application is how recordings of evoked neural activity can be used to achieve personalized VNS dosing for research subjects or patients. Second, these key results support the eCAPTURS framework (eCAPs To Unravel Responses to Stimulation) depicted in Fig. 1A, which can be used as a guide for understanding the neural and physiological effects of VNS. While the version depicted in Fig. 1A reflects the experimental set-up of this paper, in the discussion we outline how it could be adapted for other VNS settings.

## 2 Methods

### 2.1 Surgical methods

The effects of VNS on physiological and neural response were examined in six female swine (Yorkshire, (43.2 *±* 3.5) kg). Animal protocols were approved by the Institute of Animal Care and Use Committee of American Preclinical Services (Minneapolis, US). All animals were sedated with a mixture of Ketoprofen (2.0-3.5 mg kg*^−^*^1^), Tiletamine/Zolazepam (3.5-5.5 mg kg*^−^*^1^) and Xylazine (1.5-3.5 mg kg*^−^*^1^). Isoflurane (0-5%, ventilation) was used to induce anesthesia. Following intubation, the anesthesia was maintained with propofol (2-8 mg kg*^−^*^1^, i.v.). Mechanical ventilation was provided prior to VNS. During VNS, mechanical ventilation was turned off to measure the breathing response to stimulation. Normal body temperature was maintained between 38 °C and 39 °C using a heated blanket and monitored with a rectal probe thermometer. The depth of anesthesia was assessed by monitoring heart rate, blood pressure, respiration, and mandibular jaw tone.

Animals were positioned supine with both forelimbs and head extended to expose the ventral aspect of the neck. A 10 cm incision was made 2 cm right of the midline. Subcutaneous tissues were dissected and retracted to expose the underlying muscle layers. The sternomastoid muscle was bluntly separated and retracted laterally away from the surgical field. The carotid sheath was exposed and the vagus nerve was identified and isolated away from the carotid artery. The carotid artery was suspended with vessel loops and retained to the side of the surgical window with forceps.

Eight centimetres of the vagus nerve were stripped and isolated caudally of the nodose ganglion. Three cuffs (Microprobes, NC, US) were placed about 2 cm caudally to the nodose ganglion, with the most caudal cuff used for stimulation (Fig. 2). Recording was done from two cuffs (Cuff A and B), comprised of 2 ring-shaped contacts and 6 square contacts, circumferentially placed around the nerve. An inter-cuff distance of (50.0 *±* 5.0) mm was maintained between the centres of the stimulation cuff (Cuff C) and the most rostral recording cuff (Cuff A). Two cuff design variants were used for stimulation across the study cohort. Group 1 (subjects S1, S2, S3, S4, S6) used the same design as the recording cuffs. Group 2 (subject S5) used a cuff with 16 small square contacts in longitudinal pairs arranged in the same circumferential pattern around the vagus. An implantable EMG needle was inserted in the muscle left medial to the nerve for referencing the acquisition system to the tissue potential.

**Figure 2:**
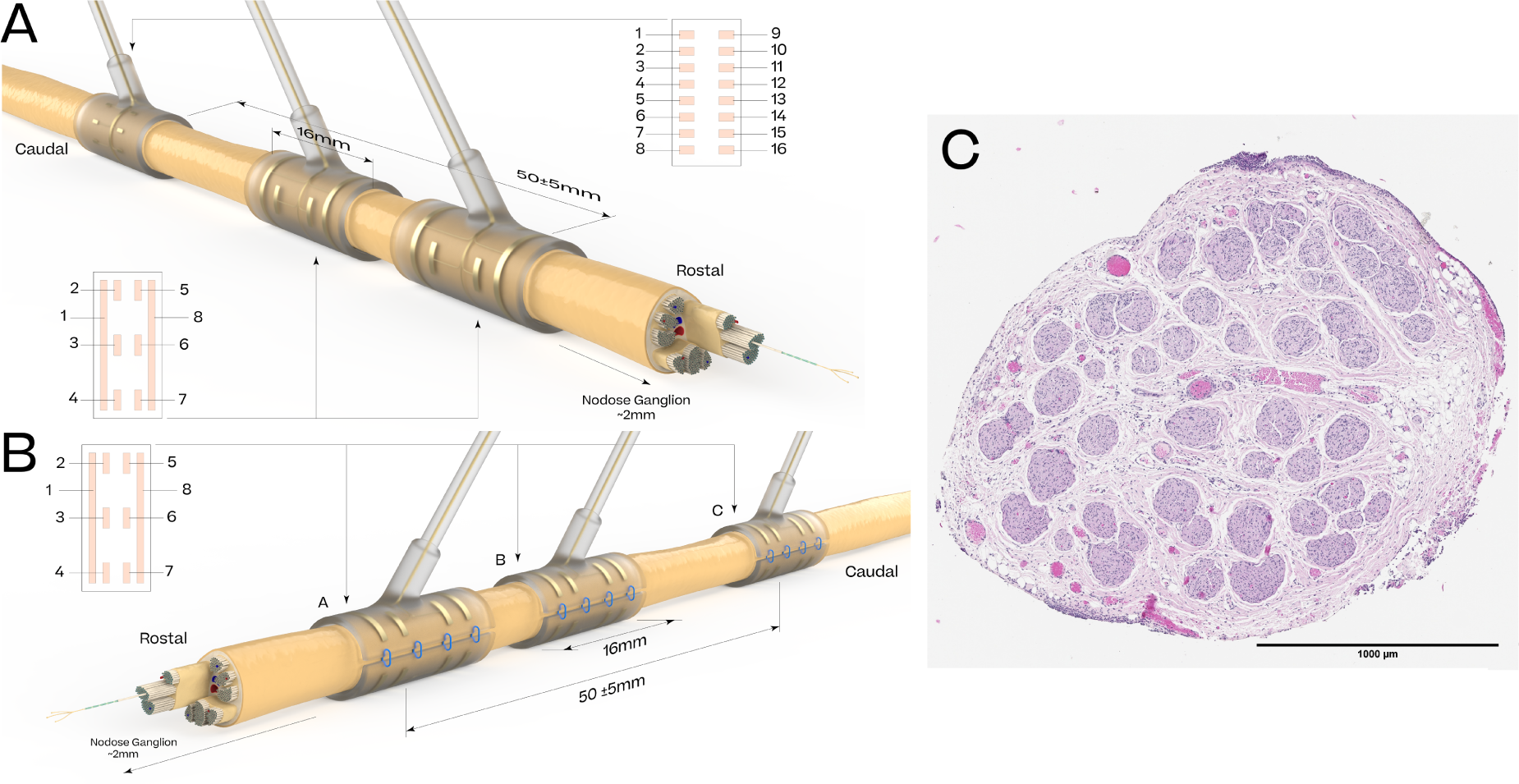
Representation of the cuff layout for Group 1 (A), Group 2 (B) subjects, and illustration of the histology of the vagus nerve (C).

At the end of the study, the swine were humanely euthanised with an intravenous injection of Euthasol (390 mg mL*^−^*^1^). Death was confirmed by an absence of pulse on the ECG and arterial blood pressure signals.

### 2.2 Custom neural interface

Cuff electrodes were connected to a custom neural interface utilising a 32-channel biopotential amplifier headstage (RHD2216, Intan Technologies) and a 16-channel bidirectional stimulator/amplifier headstage (RHS1116, Intan Technologies). The two headstages are controlled by an Artix 7 FPGA (Xlinix) sampling up to 30 kSamples/s per channel. Processes for data acquisition, data management, analysis and distribution are managed by Jetson TX-2 (Nvidia, CA, USA). This custom neural interface can be used on freely moving animals.

### 2.3 Physiology monitoring

A two-lead electrocardiogram (ECG) was used in Lead I configuration using two subcutaneous probes. Laryngeal muscle activation were monitored by electromyography (EMG) using four surface patch electrodes arranged symmetrically in two bipolar derivations, similarly to Vespa et al., 2019. Briefly, one electrode pair was placed at 6 cm above and below the laryngeal prominence and one pair was placed horizontal at half the distance between the laryngeal prominence and the medial edge of the sternocleidomastoid muscle. Both ECG and laryngeal EMG signals were amplified and digitised using a bipolar recording headstage (RHD2216, Intan Tech). Breathing was measured using a SPR-524 ultra-miniature Nylon pressure catheter (ADI, Colorado, US) and arterial blood pressure was measured with a 2F catheter implanted in the right femoral artery. Breathing and blood pressure recordings were digitised at 1 kHz (PowerLab 16/35, ADI) and then visualised using LabChart (ADI, Colorado, US).

### 2.4 Stimulation protocol

The following stimulation parameters were varied throughout this study: current (0.03-2.5 mA), frequency (1-50 Hz), pulse width (130-1000 µs), burst duration (1-10 s), and spatial location. Stimulations were either monophasic or biphasic with interphase delay, where a stimulation train consists of regularly spaced pulses of alternating polarity. Bipolar stimulation was delivered from different combinations of electrode pairs in the stimulation cuff. As a result, the depolarisation site where the eCAPs originate from (ie the current source) corresponds to either one or the other electrode, depending on the polarity. Most stimulations performed used *longitudinal* pairs of contacts, where the source and sink are located along the nerve (e.g. C2-C5, for contact labels see Fig. 2), and a small subset of stimulations used *radial* pairs of contacts, where the source and sink are located around the nerve (e.g. C2-C3). For longitudinal pairs, we use the convention that the current source corresponds to the more rostral contact for *cathodic* pulses and to the more caudal contact for *anodic* pulses. We use the following channel naming convention: two stimulation electrodes, e.g. C1-C8, with the first electrode being the most rostral of the two, and *cathodic pulse* or *anodic pulse* to indicate which electrode of the pair serves as the depolarization site.

Stimulation parameters were sequentially applied from a grid of parameter combinations. Prior to stimulation, the onsite electrophysiologist would mark when the hemodynamic indices had settled. We would apply the stimuli remotely, record the evoked neural and cardiovascular responses, and wait for the vitals to return to baseline before restarting the cycle. For stimulations affecting physiology, we would leave a minimum of 30 s between each stimulus. If the stimuli were deemed to be causing discomfort, we would not raise the charge further. If present, these would be in the form of heavy coughing or extended bradypnea or apnea. For this reason, it was not always possible to apply the same grid ranges to different subjects, when the focus was on physiological effect of VNS. This resulted in parameter sets that are only partially overlapping between subjects and made inter-subject comparisons less useful. With a focus on neural activation with short stimulation durations, however, similar sets of parameters were applied to different subjects and some direct inter-subject comparisons were possible.

### 2.5 Data processing

#### Physiological effects

R-peaks in the ECG are identified by peak finding. The *heart rate* (HR) is calculated from RR distance and interpolated by the interpolate function (method pchip) from the Pandas (team, 2020) Python package. *Heart rate change* (ΔHR) is defined by the difference of average heart rate before and shortly after the onset of stimulation. The three parameters (segment before onset of stimulation, gap time after onset, segment after gap) are chosen for each subject and parameter grid individually to optimize the model fit (measured by *R*^2^) of a linear regression model with stimulation parameters as the explanatory variables. Examples of heart rate changes in response to stimulations are given in Supplementary Material.

Breathing pressure peaks are identified by peak finding, and the *breathing rate* (BR) is calculated by forward filling. The *breathing rate change* (ΔBR) is calculated by first computing the baseline breathing rate, which is the average breathing rate in a 5 second window before stimulation. To capture bradypnea following stimulation, the minimum breathing rate is taken in a window from the onset of stimulation to 3 seconds post stimulation and the baseline breathing rate subtracted. Examples of breathing rate changes in response to stimulations are given in Supplementary Material.

We observed two different effects on neck muscles. Activation of laryngeal muscles via the recurrent laryngeal branch by efferent A*β*-fibres as described in Nicolai et al., 2020 was observed in all subjects and validated for S6 via caudal vagotomy (Supplementary Material). In this work these are termed *laryngeal twitches*, and quantified by taking the *L*_2_-norm of the muscle artefact appearing as a result in the neurograms (see Fig. 3b). We also observed a phenomenon akin to a stronger contraction of the throat muscles, characterised by an increase in the EMG signal shortly after a stimulation. We term this effect *laryngeal spasm*, and propose to quantify it using the relative increase of the *L*_1_-norm of the EMG signal one second before and after the stimulation train (Supplementary Material).

**Figure 3:**
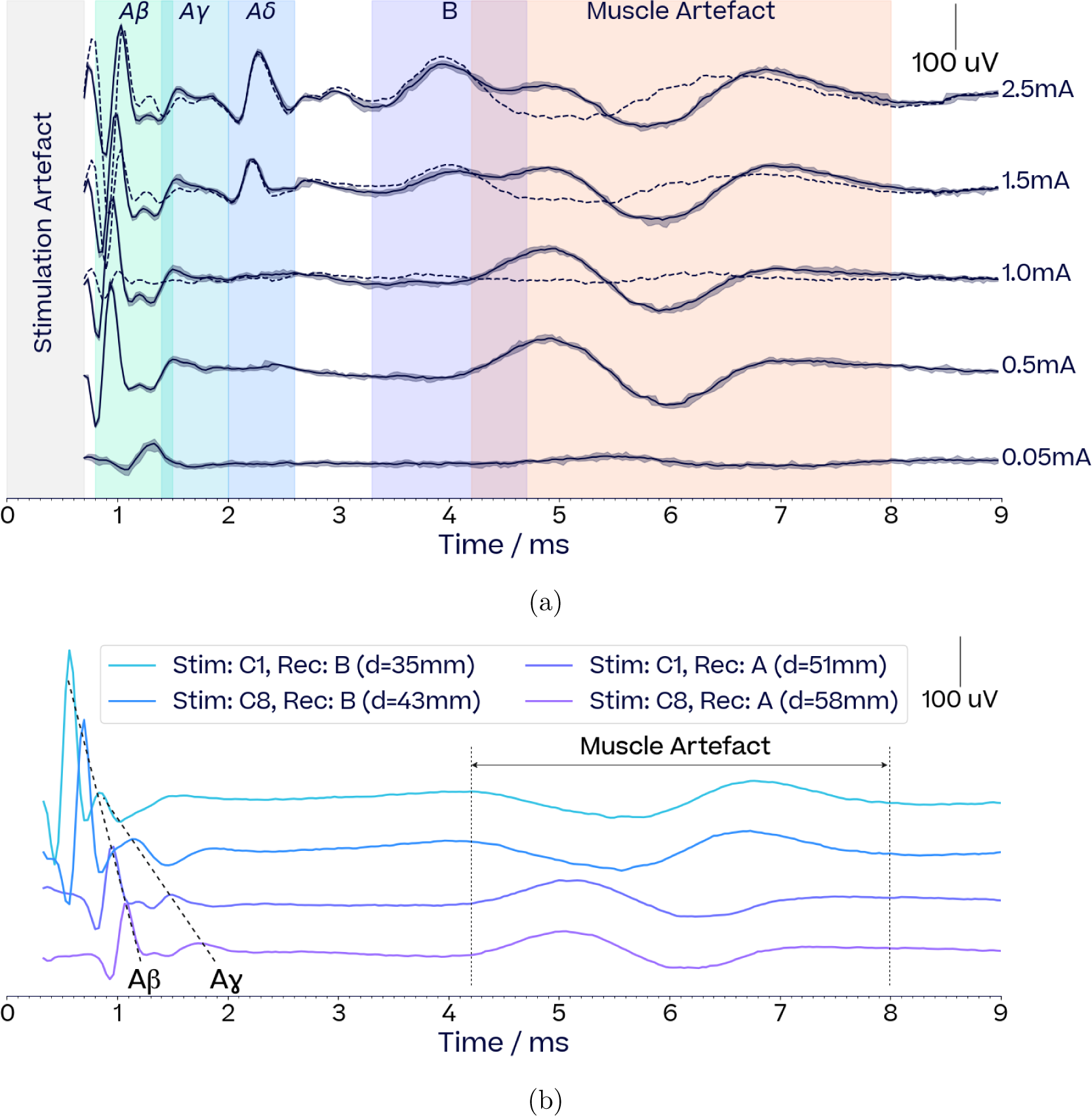
Recordings of evoked compound action potentials (eCAPs) (a) Neurograms at increasing currents (pulse width 500us, frequency 10Hz, duration 1s). Solid lines and confidence intervals respectively show the average and 5th/95th percentiles of the detrended response across the 10 pulses forming each stimulation train. For currents *>*= 1.0 mA, dotted lines show the average response after substracting the average 0.5 mA response to isolate the B-fibre eCAP from the muscle artefact. (b) Propagation of the neural signal between different stimulation sites and recording locations for A*β* and A*γ* eCAPs (current 0.25 mA, pulse width 130 µs). Cuffs A, B and C correspond to Group 1 cuff layout, as illustrated Fig. 2. Distances between the stimulation and recording site are shown in the legend. The propagation of the A*β* and A*γ* eCAPs are consistent with known conduction velocities. The muscle artefact occurring between 4-8 ms shows no propagation between recording and stimulation sites but changes polarity between recording cuffs.

#### Neural signal

Evoked compound action potentials (eCAPs) from the vagus were isolated from the recordings after identifying and removing the stimulus pulses across a stimulation train. A 5th order Butterworth high-pass filter (200 Hz) was applied to correct for any exponential decays of the stimulation pulses. The signal was averaged across all stimulation pulses and eCAPs were classified into fibre types (Erlanger-Gasser classification) based on their conduction speed (see Fig. 3a). The authenticity of neural signals was validated by looking at their propagation between different depolarisation and recording locations. This is illustrated by Fig. 3b: as the relative distance between stimulation and recording sites increases, A*β* and A*γ* eCAPs shift in proportion to their respective conduction speed, while an EMG artefact appearing around 5-7 ms does not. We did not observe any C-fibres throughout our experiments. This is not surprising considering these are typically observed for currents well above our maximal current of 2.5 mA or with other specialist preparations (Tosato et al., 2006; Yoo et al., 2013).

#### eCAP features

Activations from each fibre were computed by taking the *L*_2_-norm of the linearly detrended signal within predefined time windows. The definition of these time windows are primarily derived from each fibre’s conduction speed, but also account for virtual cathode effects (Wikswo et al., 1991) for anodic pulses, which shift the depolarisation site of the afferent eCAPs in the rostral direction as current increases (Supplementary Material). Neural thresholds were defined as the current leading to 15% of the maximal activation recorded for each fibre, and computed by linearly interpolating dose-response curves. Potential contamination of B-fibre activation computation by muscle artefacts was removed by subtracting the response curve at 0.5 mA from responses to currents above 0.5 mA (see Fig. 3a).

### 2.6 Data analysis

#### Train and pulse parameters

An important methodological assumption we make in this study is separating VNS parameters into two groups–*pulse parameters* (applied current, pulse width, and electrode location) and *train parameters* (frequency and train duration). We assume that pulse parameters determine the evoked activation pattern, while train parameters integrate the evoked neural activity into physiological responses. In the frequency range typical of VNS, train parameters have no influence on the shape of neural activations. The limited effect of train parameters on neural activation has been observed previously (for example, Chang et al. (2020)). However, in this study we systematically explore the effects of these two sets of parameters and discuss how this might inform a new method for efficient VNS parameter optimization. We routinely verify that neural activations are unaffected by train parameters by inspecting eCAP profiles generated for a range of train parameters: an example is shown in Fig. 8.

#### Alignment criterion for comparing neural activations with physiological effects in a subject

We consider the possibility of a causal connection between a neural and a physiological response to stimulation whenever the dose response curves (DRCs) agree in their points of onset and saturation (Verma et al., 2022; Blanz et al., 2022). We might term this approach the *DRC alignment criterion* for a potential causal connection between a specific fibre type activation and a specific physiological response. This is further discussed in Section 4.3.1.

#### Gaussian process models

*Gaussian process models* (Rasmussen and Williams, 2006) are used throughout the study for nonlinear regressions of response data. Nonlinear response curves and surfaces are obtained as mean functions of Gaussian processes with either squared exponential or Matern kernels. Hyperparameters are fitted by maximising the likelihood while imposing Gamma distributions as priors on parameters.

### Regression functions

A *two-exponential curve* is computed by fitting a parametric curve of the sum of exponential functions: *f* (*x*) = *a*_0_ +*a*_1_ exp(*−x/b*_1_)+*a*_2_ exp(*−x/b*_2_). The fit is obtained by minimizing the sum of residual squares over the five parameters. Similarly a *so*ftplus curve fits *f* (*x*) = *a*_0_ + *a*_1_ log(1 +exp((*x − a*_3_)*/b*, and a *sigmoid* curve *f* (*x*) = *a*_0_*/*(1 +exp(*−*(*x − a*_1_)*/b*). A *step-sigmoid* curve first fits a sigmoid curve to points with *x < c* and then a second sigmoid curve to residuals with *x > c*.

#### Assessment of regression fit

To determine the fit of nonlinear regression models or the improvement of fit when comparing two regression models we use effective degrees of freedom of the kernel matrix and an *approximate F-test* (Hastie, Tibshirani, and Friedman (2009), Sec. 5.5.1).

## 3 Results

The results are organised in four subsections: (1) the inter-patient variability of physiological responses to VNS, (2) the importance of a high-dimensional parameter search for individualized VNS dosing, (3) the effect of stimulation parameters on evoked neural activity and (4) the link between evoked neural activity and physiological responses.

### 3.1 Inter-subject variability of VNS responses

Fig. 4 compares percentages of changes in heart rate of several subjects in response to VNS for various combinations of frequencies and train durations or currents. The subjects show markedly different responses to the same stimulation parameters. In Fig. 4a, both S1 and S3 exhibit bradycardia and tachycardia in response to VNS; however, the precise parameters that trigger these effects differ between subjects, as does the strength of each effect. In Fig. 4b, S4 (purple) shows a strong increase in tachycardia with an increase in train duration from 2 to 3 s, while S3 (green) has a comparatively flat response over the whole parameter range. In Fig. 4c, S6 (purple) responds with bradycardia, while S3 (green) shows mostly tachycardia over the same range of frequencies and currents. These examples strongly suggest the need of patient specific VNS parameter adjustments to achieve defined physiological responses.

**Figure 4:**
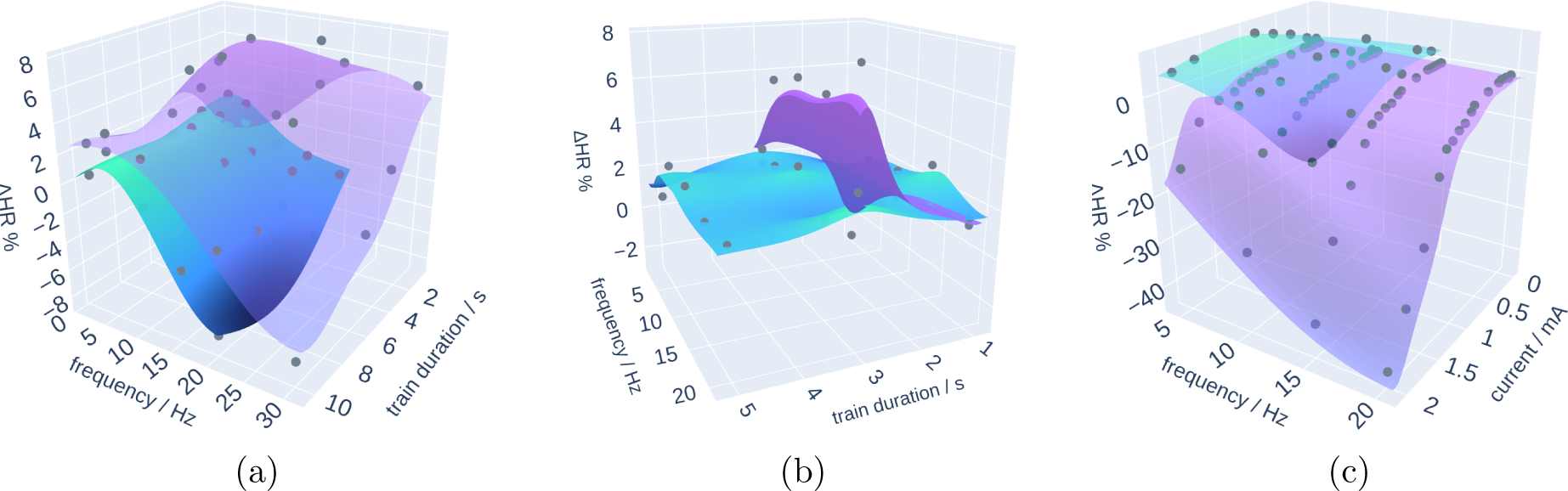
Comparison of ΔHR response in different subjects. (a) Subject S3 (purple) compared to S1 (green) (current 1.5 mA, pulse width 500 µs), (b) Subject S4 (purple) to S3 (green) (current 0.8 mA, pulse width 500 µs, (c) Subject S3 (green) compared to S6 (purple) (train duration S3 3 s, S6 5 s, pulse width 500 µs))

### 3.2 Response to vagus nerve stimulation using multiple parameters

In this section we present examples that show that in many cases optimizing physiological responses to VNS over multiple parameters is preferable to optimization over a single parameter. We explore the effect of a reduction of multiple parameters to total charge when optimizing heart rate, and the advantage of considering multiple stimulation parameters or electrode contact locations when optimizing heart rate while controlling off-target effects such as bradypnea or laryngeal spasms.

#### 3.2.1 Limitations of reducing VNS parameters to charge

Stimulation dosage in form of total charge is an important determinant of a therapeutic effect (Zaaimi, Grebe, and Wallois, 2008; North et al., 2021). In Fig. 5b a regression surface (grey) based on total charge is compared to a GP regression surface (green) based on the two separate parameters of frequency and current. This is achieved by computing the charge for each frequency-current input point and retrieving the corresponding response from the regression curve in Fig. 5a. The total charge surface explains most of the variability. One might therefore feel motivated to simplify the search for an optimal VNS dosing by combining several parameters into a single composite metric such as total charge.

**Figure 5:**
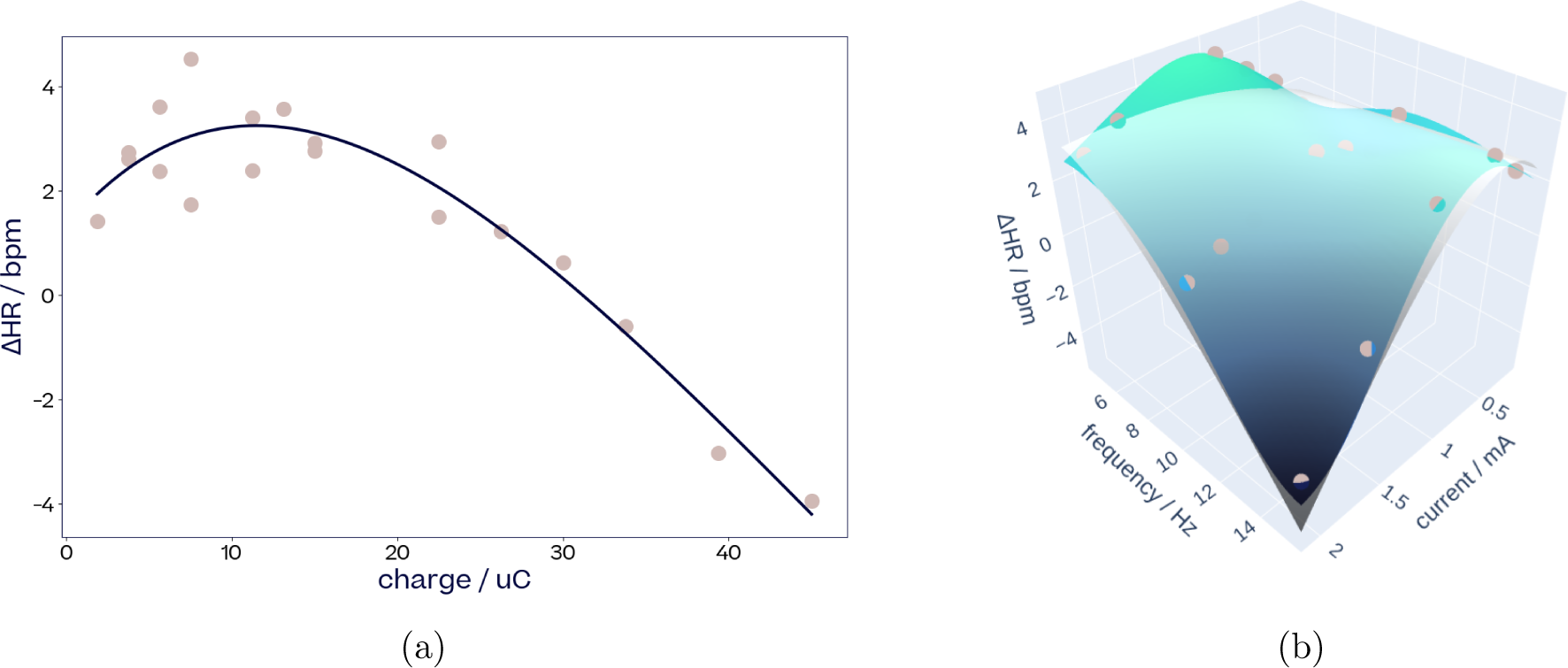
ΔHR response surface fit based on a single parameter (total charge) compared to fit based on two parameters (current and frequency). (a) Fit of a two-exponentials function to ΔHR data from subject S3 (train duration 3 s, pulse width 500 µs), this fit is transferred to (b) as grey surface. (b) Two dimensional GP fit (green) to the same data on top of a one dimensional fit (grey) from (a) transferred to two dimensions. The difference in fit is significant (F-test, *p <* 10*^−^*^11^).

However, Fig. 5 also illustrates how such a metric can lead to sub-optimal results. The GP regression surface over the full parameter space has a significantly better fit (F-test, *p <* 10*^−^*11) and explains more of the variability of the HR response. If, hypothetically, the aim was to find VNS parameters that maximise tachycardia, the charge curve would suggest an optimum around 3 bpm, while a stimulation based on the expanded current and frequency space would achieve higher tachycardia of around 5 bpm.

This example shows that simplification of the parameter space in order to accelerate parameter searches leads to sub-optimal results compared to a full dimensional search.

#### 3.2.2 VNS parameters for a trade-off between on-target and off-target responses

As on-target effects are often achieved by parameters that also evoke side effects, we explored a variety of systematic and extensive grids of VNS parameters for their effect on heart rate changes, which we consider an on-target effect, and breathing rate changes and laryngeal spasms, which we consider off-target effects.

With three examples we illustrate how a search in a two dimensional parameter space (frequency combined with train duration or frequency combined with current) leads to improved optimization results for an on-target effect while avoiding an off-target effect, compared to a search along only one parameter. Changes in heart rate and breathing rate were recorded in subjects S4 and S6 in response to frequency and train duration or current (Figs. 6a, b and Figs. 6d, e), while changes in heart rate and laryngeal spasms were recorded in subject S3 in response to frequency and train duration (Figs. 6g, h).

**Figure 6:**
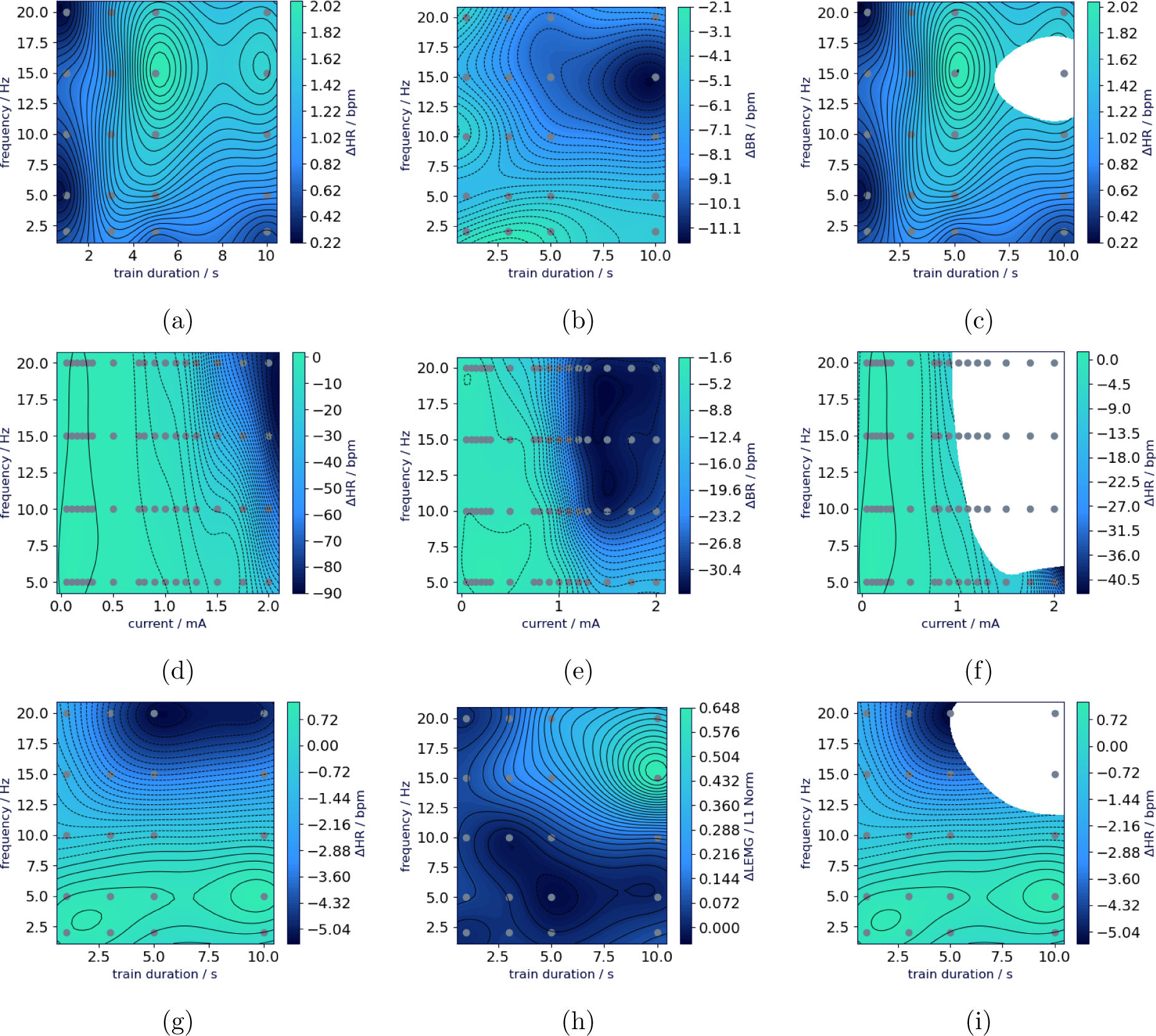
On-target ΔHR restricted by off-targets, ΔBR and lEMG. Subject S4 (a) ΔHR, (b) ΔBR and (c) the frequency-train duration space available (non-blank) for ΔHR adjustment when ΔBR *< −*10 bpm should be avoided (current 0.25 mA, pulse width 250 µs). Subject S6 (d) ΔHR, (e) ΔBR and (f) the frequency-current space available (non-blank) for ΔHR adjustment when ΔBR *< −*17 bpm should be avoided (train duration 5 s, pulse width 250 µs). Subject S3 (g) ΔHR, (h) ΔlEMG and (i) the frequency-train duration space available (non-blank) for ΔHR adjustment when ΔlEMG *>* 0.3 of maximum activation should be avoided (current 1.5 µA, pulse width 250 µs).

For subject S4, there is an optimal frequency around 15 Hz, where the train duration can be adjusted to induce bradycardia and avoid bradypnea (Fig. 6c). For subject S6, although bradycardia and bradypnea regions largely overlap (Figs. 6d, e), a corner region in the frequency-current space induces bradycardia with less bradypnea than other stimulation combinations (Fig. 6f). For subject S3, bradycardia (Fig. 6g) with reduced laryngeal spasms (Fig. 6h) is possible if frequency is restricted to about 5 Hz (Fig. 6i).

#### 3.2.3 Contact location affects the trade-off between on-target and off-target responses

VNS recordings for subject S4 illustrate how contact location affects the trade-off between tachycardia (on-target) and strong bradypnea (off-target). Monophasic stimulations at 10 and 20 Hz, and 0.25 and 0.8 mA were applied with both polarities from three longitudinal contact pairs at different radial locations on the vagus (see Methods, Fig 2). Tachycardia and bradypnea were induced from all contact pair. However, for some contact pairs increase in heart rate came at the price of stronger bradypnea than for others. Fig. 7 demonstrates that a gain in heart rate is associated with a significantly lower bradypnea for stimulations from C2-C5-cathodic than from C4-C7-cathodic (ANOVA, *p <* 0.04) and C2-C5-anodic (*p <* 0.05). This illustrates the potential of stimulating from well chosen spatial locations for optimizing a trade-off between on- and off-target responses.

**Figure 7:**
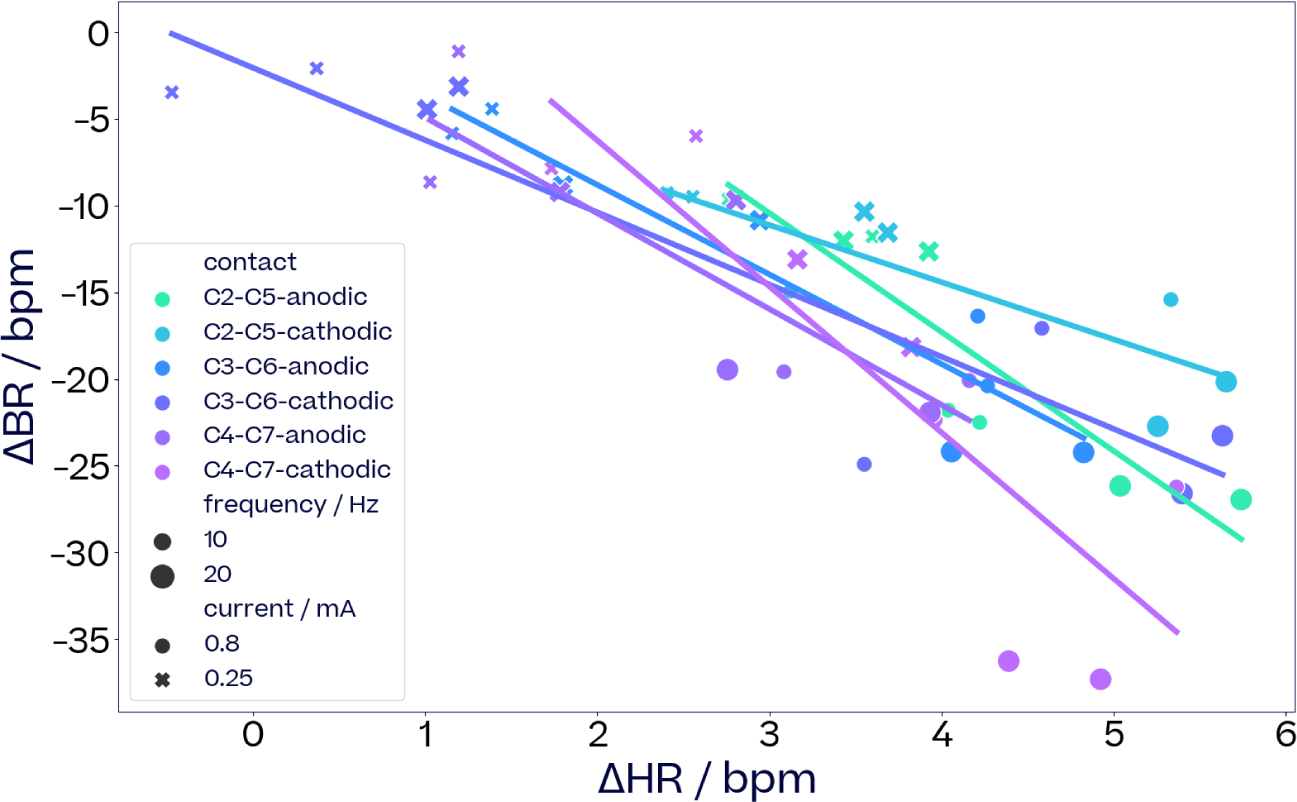
Relationship of ΔHR to ΔBR dependent on electrode location. Effect on HR and BR for six different stimulation locations (three electrode pairs, two polarities for each) pulse width was 500 µs with train durations of 3 s and 5 s, current is in mA and frequency in Hz. The train duration had no statistical effect on the relationship and is therefore not specifically labelled in the graphs. The overall location effect is significant (ANOVA, *p <* 0.0026). There is a significant difference in the slopes of C2-C5-cathodic vs C4-C7-cathodic (ANOVA, *p <* 0.04) and C2-C5-cathodic vs C2-C5-anodic (*p <* 0.05).

### 3.3 Effect of stimulation parameters on eCAPs

In this section we investigate the relationship between stimulation parameters and the evoked neural activity. The first observation, which simplified the rest of the exploration, is that frequency and train duration (collectively called train parameters) have little influence on the amount and shape of evoked compound action potentials within a frequency range of 2-50 Hz (MAPE: 0.05% *±* 0.02%). This range includes frequencies used in approved VNS therapies (2-30Hz). When current, pulse width, and electrode location remain fixed, neurograms are indistinguishable across different combinations of train parameters (see Fig. 8). This observation allows to separate the contribution of pulse and train parameters to neural and physiological responses to VNS throughout this study, as illustrated Fig. 1.

**Figure 8:**
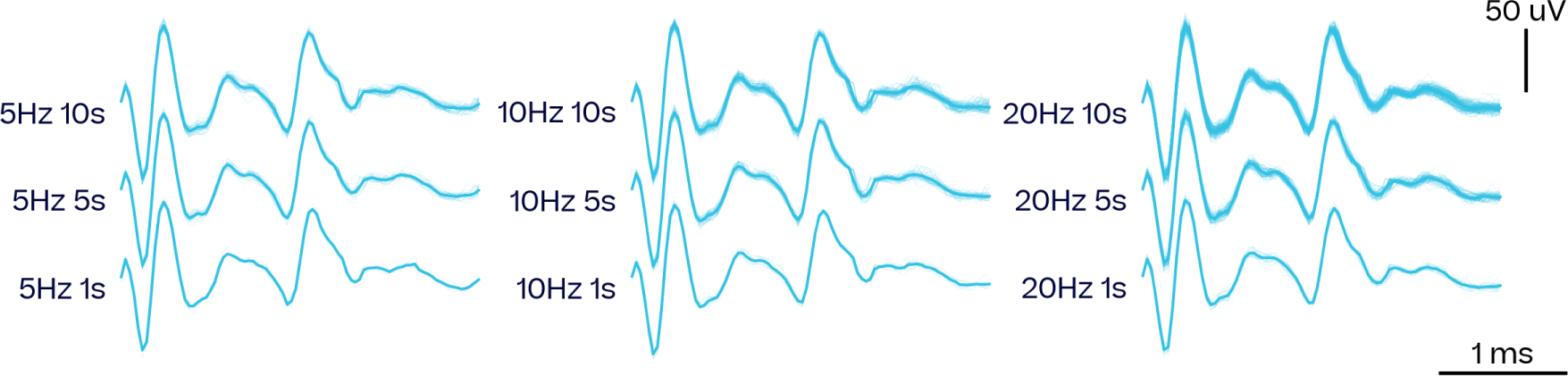
Invariance of evoked neural activity with respect to train parameters. Evoked neural activity recorded across different frequencies and train durations for fixed current and pulse width (1.5 mA, 500 µs) from subject S3. For each train, both individual pulses (thin lines) and averaged response (thick line) are plotted. Mean absolute percentage error (MAPE) of the average response with respect to the 10 Hz, 5 s response: 0.05% *±* 0.02%.

#### Fibre recruitment depends on pulse parameters

In contrast, variations in applied current, pulse width and electrode location (collectively called pulse parameters) lead to different fibre activation profiles. Current is the main driver of fibre recruitment. Fibre activation increases with current until it reaches saturation, giving rise to a sigmoidal dose-response curve (DRC). DRCs are characterised by the fibre activation (or neural) threshold and the saturation threshold, and these vary depending on the fibre type, pulse width and electrode location (see Fig. 9a for DRCs in a single subject with pulse widths from 130 µs to 500 µs, from four different stimulation locations).

**Figure 9:**
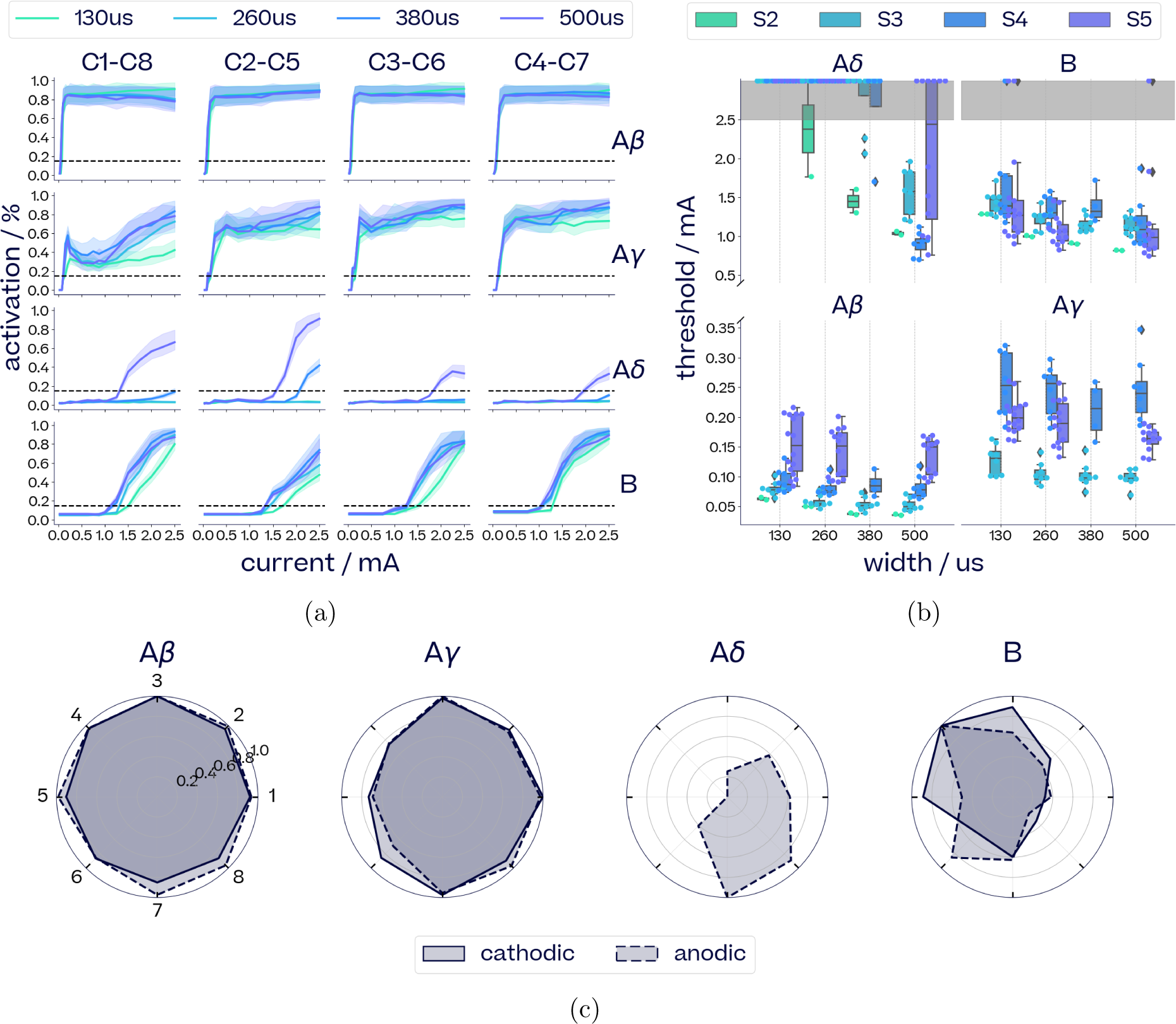
Effect of pulse parameters on fibre recruitment. (a) Dose response curves, normalised by the maximum found per fibre for subject S3. (b) Distribution of neural thresholds across different subjects and pulse widths. The maximum amplitude was 2.5 mA, and points at 3.0 mA signal that the threshold was not reached by 2.5 mA. Pulse width was found statistically significant across all fibre types (ANOVA F-test: *p <* 5.7e*−*17, *F* = 76.6) and for each fibre type (A*β*: *p <* 8.5e*−*3, *F* = 7.2; A*γ*: *p <* 7.5e*−*4, *F* = 12.1; A*δ*: *p <* 6.8e*−*15, *F* = 94.5; B: *p <* 1.4e*−*3, *F* = 10.8). (c) Maximal fibre activation reached from each location in S5, normalised by the highest activation reached across locations for each polarity. For these monophasic stimulations, the first (second) contact in each pair is the cathode in cathodic (anodic) pulses.

Fig. 9b shows the effect of pulse width on the distribution of neural thresholds for each fibre type across four subjects. For graphical display purposes, a nominal threshold of 3 mA was assigned when no activation was achieved over the current range to 2.5 mA. For all four subjects, pulse width has a statistically significant effect on neural threshold across all fibres (ANOVA F-test: *p <* 5.65e*−*17, *F* = 76.6). The effect of pulse width on the neural threshold is particularly pronounced in the case of A*δ*-fibres.

Spatial selectivity for fibre activation was observed in subject S5, in which a higher density electrode cuff was used. Fig. 9c shows the maximum activation reached for each fibre across eight longitudinal electrode pairs for current and width in the range 0.03-2.5 mA and 130-500 µs respectively. While the faster and larger A*β*-fibres are generally activated to the same extent regardless of stimulation location, smaller A*γ*-, A*δ*- and B-fibres show distinctive spatially selective activation patterns.

#### Inter- and intra-subect variability of nerve responses

We observe a wide distribution of neural thresholds across subjects (Fig. 9b). For each subject, the variation of the neural threshold of a given fibre across stimulation locations is represented by the spread of each box. The generally broader ranges of neural thresholds observed in subject S5 correspond to higher spatial selectivity owing to a different electrode layout.

### 3.4 Evoked compound action potentials as biomarkers of VNS responses

Physiological responses to VNS are influenced, on one hand, by the set of fibres activated at each stimulation pulse, or eCAPs, on the other hand, by the frequency, train duration and other time domain parameters of the stimulation. In this section we explore both sides: connections between eCAPs and physiological responses, and how train parameters contribute to the integration of eCAPs into different physiological effects. We also observe that the functional relationship between eCAPs and physiology might be considered simpler than that between VNS parameters and physiology.

#### 3.4.1 Alignment of eCAPs and physiological effects

The alignment of neural and physiological response curves can be used to highlight causal connections between physiological responses and eCAPs (Verma et al., 2022; Blanz et al., 2022).

Fig. 10A shows breathing rate changes in Subject S5 depending on pulse numbers. Linear models are fitted for each current and their slopes are shown in Fig. 10B, alongside the dose response curves of A*β*-, A*γ*- and B-fibres in Fig. 10C. As the number of pulses (train duration *∗* frequency) increases, the breathing rate decreases for all currents tested (Fig. 10A). The slope of the decrease in breathing rate is steeper at 0.75 mA compared to 0.25 mA and plateaus for higher currents (Fig. 10B). Out of the three neural fibres recruited throughout these stimulations (Fig. 10C), the DRC of A*γ*-fibre seems to show the best alignment with the change in breathing rate as it is partly activated at 0.25 mA and saturated beyond 0.75 mA. Conversely, A*β*-fibres are already nearly saturated at 0.25 mA and B-fibre activation gradually increases up to 1.5 mA.

**Figure 10:**
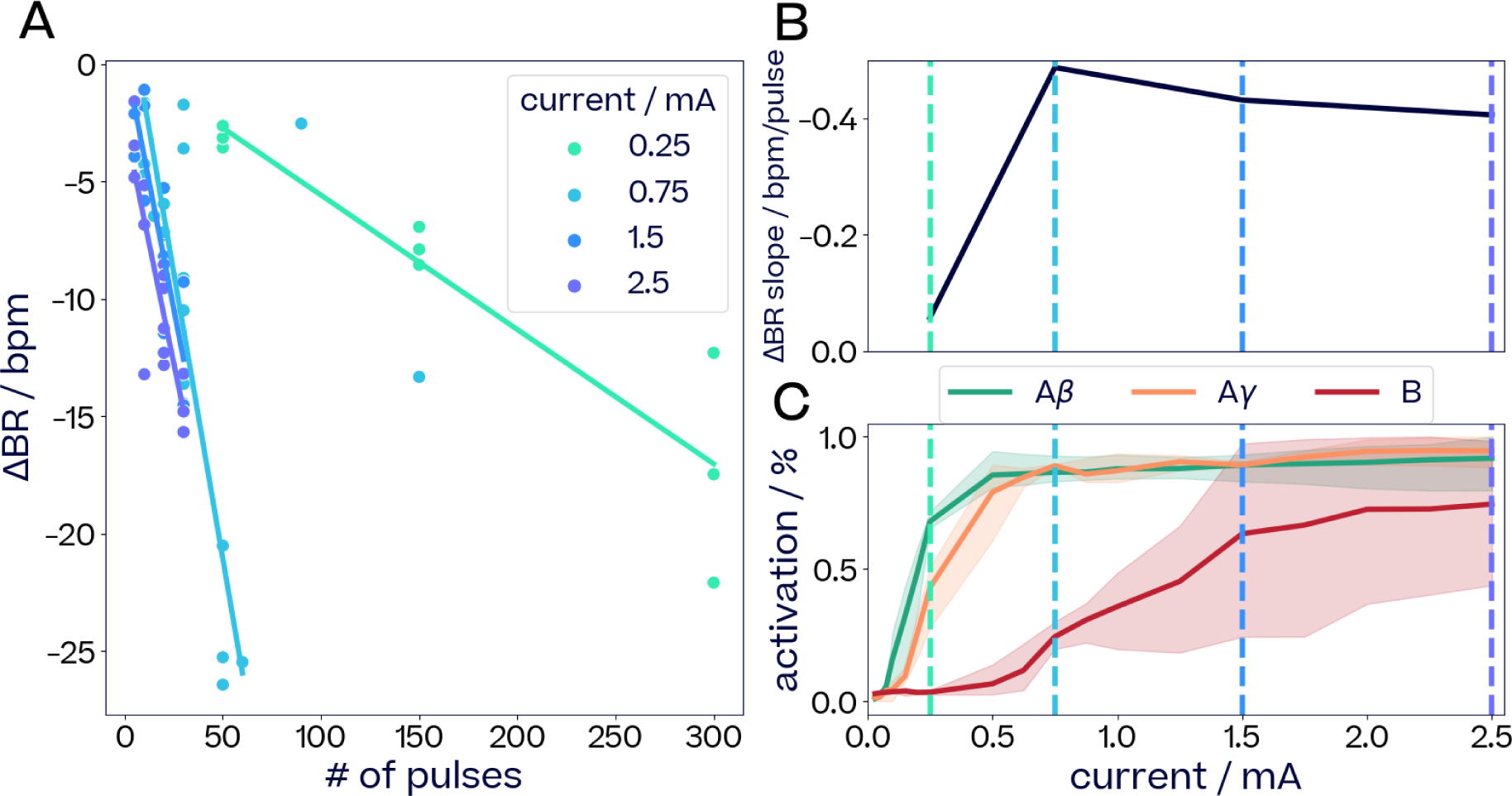
Comparison of breathing rate change with activation of neural fibres for subject S5. (A) Changes in breathing rate with respect to number of pulses (ie frequency *∗* duration) for different current amplitudes (pulse width 500 µs). Each stimulation was repeated across three different longitudinal pairs. Linear models are fitted for each current; (B) slope of linear models from (A) with respect to current amplitude; (C) dose-response curve of A*β*-, A*γ*- and B-fibres at 500 µs averaged stimulation locations, relative to the maximum activation recorded at this pulse width. Vertical lines indicate currents 0.25 mA, 0.75 mA, 1.5 mA and 2.5 mA.

For subject S6, we compare changes in both heart rate and breathing rate to the activation of A*β*-, A*γ*-, A*γ*_2_- and B-fibres in Figure 11. For increasing currents, we report ΔHR and ΔBR for frequencies between 5 Hz and 20 Hz with 5 Hz increments (top row). Biphasic pulses with an interphase delay were applied, so that a single stimulation train consists of regularly spaced pulses with alternating polarity. Fibre activations are shown for both polarities (bottom row). Mild bradycardia is induced at 0.8 mA (solid brown vertical line), which aligns with the neural threshold of B-fibres for cathodic pulses. Note that at this current, anodic pulses do not trigger any B-fibre eCAPs. As such, B-fibres are activated by every other pulse of the stimulation train. Beyond 1.2 mA, anodic pulses begin to trigger B-fibre eCAPs as well, essentially doubling the frequency of B-fibre eCAPs per stimulation train (dotted brown vertical line). This aligns with stronger bradycardia above 1.2 mA. For higher currents, both anodic-triggered B-fibre activation and bradycardia continue to increase.

**Figure 11:**
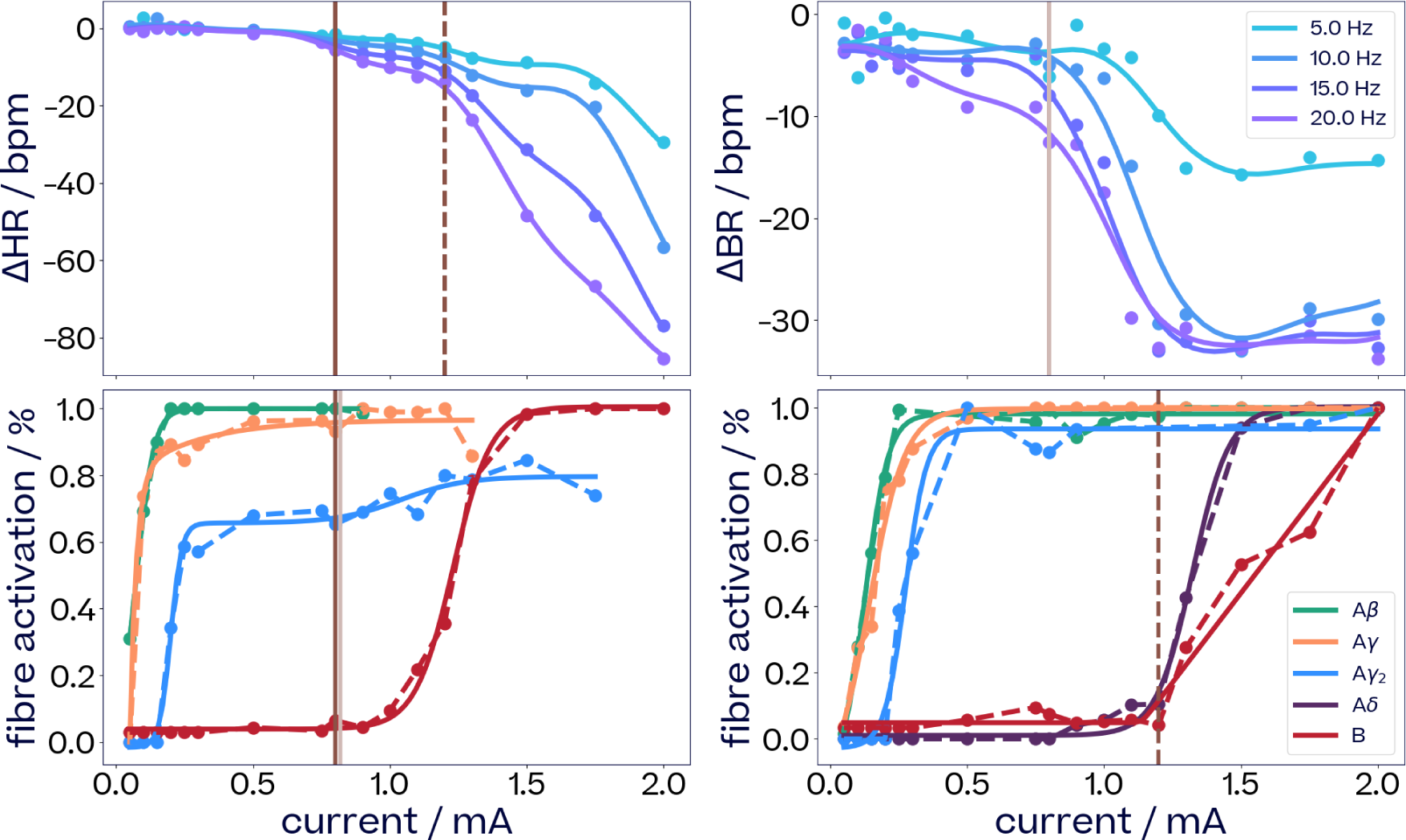
Comparison of dose-response curves of HR and BR change and neural fibre activations for subject S6. Top row: change in heart and breathing rate in response to current and frequency changes of stimuli (pulse width 500 µs, train duration 5 s). The curves are slices through a GP regression surface over the 2D input space. Vertical lines (brown) for ΔHR indicate the onset of mild bradycardia at 0.8 mA (solid) and of a stronger one at 1.2 mA (dotted). A vertical line (beige) for ΔBR indicates the onset of strong bradypnea at 0.8 mA. Bottom row: activation levels for various fibre types in response to current changes (frequency is not shown since it had no effect on eCAPs). A brown vertical line at 0.8 mA indicates the onset of B-fibre activation for cathodic pulses in the left hand plot, a beige vertical line at the same current (slightly offset for visibility) indicates the onset of a secondary A*γ*_2_-fibre activation. Similarly, a dotted brown vertical line at 1.2 mA for anodic pulses in the right hand plot indicates the onset of B-fibre activation. Data points that fell below 0.9 of the maximum after a sequence reached its maximum have been removed as outliers. Top panels contain response curve fits from slices through a GP surface. The left hand bottom panel shows fits of a sigmoid (A*β*, B), a two-exponentials (A*γ*), and a step-sigmoid (A*γ*_2_) curve. The right hand bottom panel shows fits of sigmoid curves except a softplus curve for B-fibre activation.

Mild bradypnea (Figure 11, upper right panel) is observed at low currents and seems to be related to A*γ*- and A*γ*_2_-fibre activation. Strong bradypnea (Figure 11, upper right panel, beige line at 0.8 mA) aligns with an increase in A*γ*_2_-fibre activation for cathodic pulses (Figure 11, lower left panel, beige line). As cathodic A*γ*_2_-fibre activation reaches saturation around 1.2 mA, strong bradypnea levels off.

Fig. 12 shows changes in heart rate and fibre activations for subject S3. Contrary to Fig. 11 for S6 which shows the result of biphasic stimulations with cathodic and anodic pulses interleaved, Fig. 12 shows the result of monophasic stimulations with only cathodic (left) or anodic pulses (right) and their respective HR effects. In contrast to subject S6, S3 showed pronounced tachycardia for lower currents and frequencies and bradycardia only for higher currents and frequencies. Similar to Fig. 11, B-fibre activations (lower panels) seem to follow the DRC alignment criterion with the bradycardia effects (upper panels). On the other hand, the tachycardia occurs alongside A*β*-, A*β*_2_- and A*γ*-fibre activations at low currents. For 10 Hz stimulations, it gradually increases in intensity up to 1.0 mA, which is mirrored to some extent by the activation of A*γ*-fibres.

**Figure 12:**
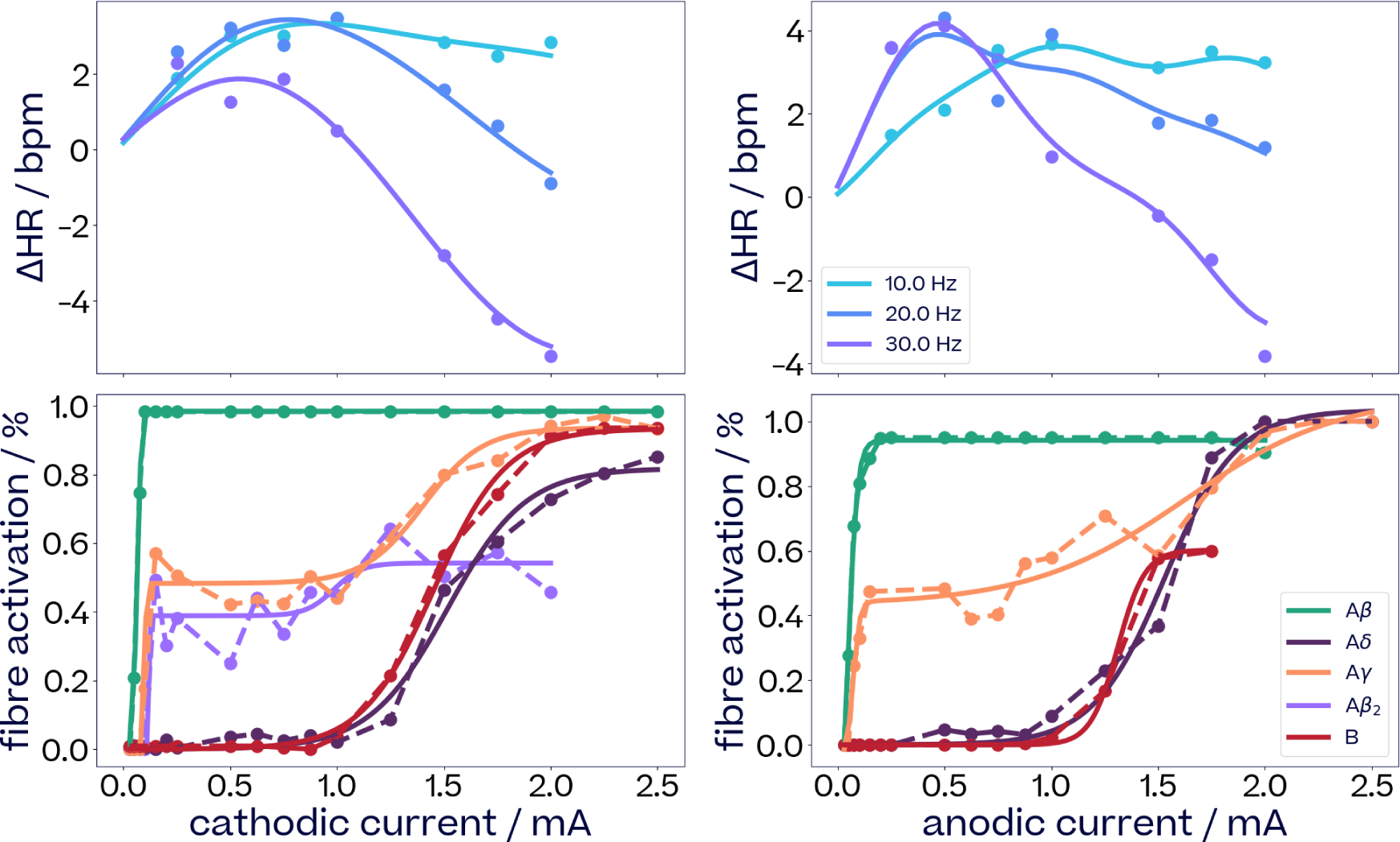
Comparison of dose-response curves of HR change and neural fibre activations for subject S3. Responses to cathodic (left) and anodic (right) stimulations. Top row: change in heart in response to current and frequency changes of stimuli (pulse width 500 µs, train duration 5 s). The curves are slices through a GP regression surface over the 2D input space. Bottom row: activation levels for various fibre types in response to changes in current (since frequency changes had no effect on eCAPs they are not shown). Data points that fell below 0.9 of the maximum after a sequence reached maximum have been removed as outliers. Top panels contain response curve fits from slices through a GP surface. The bottom panels show fits of a sigmoid (A*β*, A*δ*, B) a step-sigmoid (A*β*_2_, A*γ*) curves.

#### 3.4.2 Train parameters determine integration of eCAPs resulting in physiological effects

For a fixed set of pulse parameters, frequency and train duration determine how the neural activity evoked by each stimulation pulse integrates into physiological effects. In Figure 11, when only cathodic pulses lead to B-fibre recruitment (brown line), HR decreases gradually as frequency increases from 5 Hz to 20 Hz. When both cathodic and anodic pules lead to B-fibre recruitment (dotted brown line), there is a sharp increase in the rate of frequency-driven HR decrease. BR also decreases gradually with increasing frequency for currents above 0.5 mA, but seems to reach a plateau for frequencies above 10 Hz at high currents.

A striking effect of Fig. 12 is the difference in the integration of fibre activations that induced tachycardia from those that induced bradycardia. At currents below 1.0 mA, tachycardia is observed across all frequencies. Conversely, for currents above 1.0 mA, tachycardia is still observed at 10 Hz but transitions to bradycardia as frequency increases to 30 Hz.

#### 3.4.3 eCAPs simplify the mapping to physiology compared to stimulation parameters

Fig. 13 shows the amplitude of laryngeal contractions with increasing current and A*β*-fibre activation across various longitudinal and radial electrode pairs in S2. The relationship between laryngeal contractions and pulse parameters is non-linear and varies greatly across electrode pairs. When plotting this effect against A*β*-fibre activation, however, the relationship becomes linear regardless of current amplitude and stimulation location.

**Figure 13:**
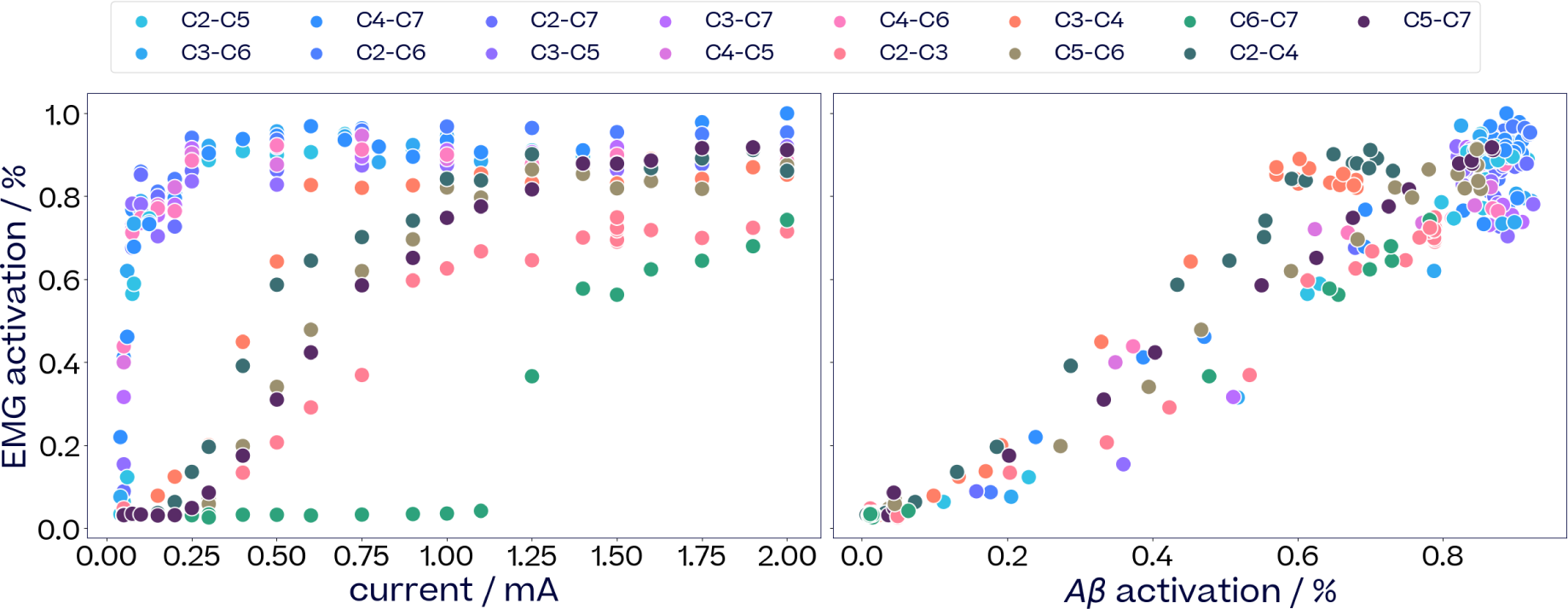
Amplitude of VNS-induced laryngeal contractions in S2. Plotted against current amplitude (left) and activation A*β*-fibres (right) across different radial and longitudinal stimulation locations. In radial pairs, the stimulation electrodes are located closer to each other on the nerve, so more current is needed to achieve depolarization of nerve fibres. A*β*-fibre activations are normalised based on the maximum A*β*-fibre activation recorded across all pairs.

## 4 Discussion

In this section, we discuss the implications of the findings presented in this work for gaining a more comprehensive understanding of the mechanisms of action of VNS. We then present a novel VNS dosing procedure which leverages these insights to improve the efficacy and accuracy of VNS therapies. Finally, we address the limitations of this study.

### 4.1 Discussion of results

#### 4.1.1 Variation between subjects

We observed considerable inter-subject variability and some intra-subject variability in physiological and neural responses to VNS (Fig. 4). At low charge dosages, most subjects exhibited tachycardia, but as charge increased, the tachycardia transitioned into bradycardia. Subject S6 (Fig. 4c), however, exhibited bradycardia in response to the whole range of VNS frequencies and currents. This was likely due to the high base heart rate (around 160 bpm) maintained throughout the experiment. Meanwhile, subject S4 only exhibited tachycardia for the range of charges that were deemed safe. Additionally, Fig. 6 illustrate how the most desirable VNS parameters vary greatly between subjects, highlighting the importance of personalized dosing in VNS therapies.

The variability in evoked neural response can best be seen in Fig. 9b, which compares the neural responses of four subjects S2 to S5. For each subject (color), the spread of each box represents the variation of neural thresholds for different stimulation locations. Notice that not only did activation thresholds of fibre types differ between subjects, but also their variation over stimulation locations: some subjects showed very consistent activation thresholds independent of electrode contact, while for others thresholds differed considerably from contact to contact.

The high inter-patient variability observed in both physiological and neural responses to VNS suggests that personalized VNS dosing procedures are critical to the efficiency of VNS therapies.

#### 4.1.2 Multiple parameters for optimal VNS dosing

One of the main challenges when using titration to optimize VNS parameters is that side effects can prevent one from reaching therapeutically effective regions of the parameter space. This is illustrated by examples in Fig. 6, where the region of the parameter space with desirable on-target effects often overlaps a region with undesirable off-target effects, an exclusion zone blanked out in white. However, if multiple parameters of a higher dimensional space are considered, a narrow region of satisfactory on-target effects avoiding unacceptable off-target effects can often be found just outside the boundary of the exclusion zone. If one simply titrated current or train duration along a fixed frequency such as 10 Hz, the best solution for maximum HR reduction with minimum side effects would be inferior to a solution found when one varies frequency in addition to current or train duration. Figs. 6c, f, and i depict this for frequencies of 15, 5, and 20 Hz, respectively.

In the examples of Fig. 6 we only investigated on- and off-target optimization over two parameter dimensions (frequency-current, and frequency-train duration), which constitute low dimensional slices through a higher-dimensional parameter space. It stands to reason that searching over more parameters would improve the likelihood of finding satisfactory VNS parameters further.

Another dimension of VNS parameters is added through a choice of electrode locations as in section 3.2.3. Fig. 7 shows the relationship between ΔHR, considered an on-target effect, and ΔBR, considered an off-target effect, for six different contact pairs of a bipolar stimulation. This relationship changed with contact pairs, some showing a higher gain in ΔHR for a smaller amount of ΔBR than others. Our dosing method introduced in Section 4.2.1 could exploit such dependencies for an optimal trade-off between on- and off-target effects by choosing the right contacts in addition to optimizing other VNS parameters.

The size of the parameter space can be reduced by a combination of parameters into a single measure for the strength of a stimulus, for example, into the total charge delivered by a pulse train as in Zaaimi, Grebe, and Wallois, 2008. However, in section 3.2.1 we compared maximizing tachycardia over charge to maximizing over a wider parameter space. In the example of Fig. 5 more of the variability in the responses is explained by a multi-dimensional space suggesting that a search over multiple parameters is preferable to a search over a single combined measure.

Overall, these examples show that considering as many dimensions of the parameter space in VNS as possible increases the opportunity to find a parameter set with a favourable balance between on- and off-target effects. This comes at a price, however: with every additional parameter the search space grows drastically and VNS dosing becomes increasingly difficult and time consuming. As we describe in Section 4.2.1, we argue that measurements of evoked neural activity allow to alleviate this issue and perform thorough searches of the parameter space quickly and safely.

#### 4.1.3 Selective eCAP recruitment via pulse parameters

Selective VNS (sVNS) is a promising way to improve the precision of VNS therapies (Fitchett, Mastitskaya, and Aristovich, 2021) by targetting some fibres or fibre types while avoiding the activation of others. The stimulations we applied in this study were restricted to square pulse shapes. Hence, fibre-selective stimulation depended on variation in pulse current, width or stimulation location (including polarity). Here we discuss the extend to which these stimulation parameters can achieve selective eCAP recruitment.

Throughout our experiments, pulse width had a significant effect on the evoked neural activity: wider pulses were associated with lower activation thresholds for currents across all fibres (Fig. 9b). This follows from the strength-duration curve of activation thresholds relating pulse widths to currents (Irnich, 2010). This effect enabled fibre-selective activation in the case of A*δ*-fibres, which were almost never recruited for pulse widths below 260 µs but consistently activated above 500 µs. The effect of pulse width on A*β*-, A*γ*- and B-fibre activation was less pronounced. The activation threshold for fast A-fibres remained much lower than that for B-fibres regardless of pulse width.

Spatial sVNS exploits the phenomenon that different fibres can be activated from different contact locations. Stimulation in subject S5 demonstrates the potential for selective activation of either A*δ*-or B-fibres (Fig. 9c) depending on contact location. Anodic stimulations from contacts 7 and 8 recruited A*δ*-fibres and little to no B-fibres, while contacts 3, 4 and 5 recruited B-fibres and no A*δ*-fibres for both polarities. Activation of A*β*- and A*γ*-fibres could only be reduced to around 80% of their maximal activation be choosing specific spatial locations. A*β*- and A*γ*-fibres are the largest myelinated fibre types and as such they seem to be activated more uniformly across electrodes. Consequently, it might be difficult to suppress A*β*- and A*γ*-fibre activation while activating other fibre types based on current and electrode location alone.

These findings suggest that, although the simulation framework considered in this work allow for some selectivity over A*δ*- and B-fibres, lower-threshold fibres such as A*β*- and A*γ*-fibres are not easily mitigated during high intensity stimulations. Nevertheless, further eCAP selectivity might be possible by using more complex pulse shapes, as discussed in Section 4.3.7.

#### 4.1.4 Selective integration of eCAPs via train parameters

In selective VNS pulse parameters such as contact location or pulse shape are chosen in order to target specific fibre types within the nerve and consequently induce specific downstream physiological effects. We observed little if any influence of train parameters on evoked neural activity in this study. Nevertheless, physiological selectivity of stimulations can be achieved via a judicious choice not only of pulse but also train parameters, an effect we term *selective integration* of eCAPs.

Prime examples of selective integration of eCAPs are seen in both cathodic and anodic stimulations of Fig. 12. VN stimulations with the same pulse parameters and hence the same eCAP profile showed very different effects on HR depending on frequency, from causing tachyto causing bradycardia. This drastic and qualitative change in physiological response can only be attributed to differential or selective integration effects of the very same eCAP pattern through a change in train parameters. We hypothesise that at low frequencies B-fibre do not exert their bradycardia effect, while faster A-fibres already exert their tachycardia effect. Meanwhile, increasing integration levels with higher frequencies lead B-fibres to exert their bradycardia effect, increasingly overpowering any underlying tachycardia effect. A similar effect has been found in mice in McAllen et al. (2018). Similarly, all stimulations in Fig. 6g, h induced the same evoked neural activity, since only train parameters were changed throughout the grid. However, a low train duration and high frequency mitigates laryngeal spasms while maintaining a strong bradycardia effect.

Consequently, systematic exploitation of selective integration of eCAPs via train parameters provides an interesting avenue to targeting physiological effects in addition to selectivity at the eCAPs level via pulse parameters.

#### 4.1.5 Alignment between recorded eCAPs and physiological effects

Although a formal causal analysis of the link between eCAPs and physiological effects is not applicable, as discussed further in Section 4.3.1, alignment of neural with physiological DRCs might be suggestive of connections between specific fibre types and physiological effects such as bradypnea, bradycardia and tachycardia.

Previous studies concluded that bradypnea might be caused by A-fibre afferents from slowly adapting pulmonary receptors in the lungs (Kubin et al., 2006), such as A*γ*-fibres. Our data suggest a similar mechanism. The bradypnea observed for subject S5 (Fig. 10A) seemed to be driven by train parameters (number of pulses) through a linear dependency. The slopes of these dependencies are shown in Fig. 10B as DRC over current. This curve aligns most closely with the activation of A*γ*-fibres in Fig. 10C.

For subject S6, mild bradypnea was observed at low currents, alongside the activation of A*γ*- and A*γ*_2_-fibres (Fig. 11). A second increase in activation of cathodic A*γ*_2_-fibres aligns with a second, stronger increase in bradypnea. Such a second raise after a first activation plateau might be attributed to the activation of an additional group of fascicles of the same type within the nerve. This effect is discussed further in Sec. 4.1.6.

Efferent B-fibres have previously been found to be the primary driver of cardioinhibitory effects (Qing et al., 2018). This is in line with our experimental results. In S6, bradycardia effects closely follow the activation of B fibres in both cathodic and anodic pulses (Fig. 11). In S3 (Fig. 12), tachycardia is offset by bradycardia from 1.0 mA for both cathodic and anodic stimulations, which aligns with the neural threshold of B-fibres. Tachycardia responses occur as a result of a decrease in neural parasympathetic drive caused by the activation of afferent sensory fibres of the VN (Yamakawa et al., 2014; Ardell et al., 2017). This is reflected in our experimental data: in S3 (Fig. 12), tachycardia appears alongside fast A-fibre activation at currents below 1.0 mA, and, at 10 Hz, increases for currents up to 1.0 mA in alignment with A*γ*-fibres.

#### 4.1.6 Refined estimation of evoked neural activity via multiple recording contacts

Throughout our experiments, activation of neural fibres were recorded from multiple locations. The amplitude of recorded eCAPs varied depending on the location and shape of the recording contacts. The variation is indicated with confidence bounds on the dose response curves. For example, Fig. 9a shows that activation of neural fibre could vary by 40% across recording electrodes in subject S3.

We hypothesize that this variability provides valuable information about the underlying evoked neural activity: eCAPs from a given fascicle will be larger from electrodes placed in their vicinity. The set of activation recorded from multiple local contacts therefore provides a sort of *activation signature* for every eCAP that can be used to characterise it further than via its propagation velocity. This is of importance since different fascicles within the vagus can share the same fibre type and propagation velocity but lead to different physiological effects. Isolating their respective eCAPs from the recordings based on their activation signature is therefore crucial to model the link between evoked neural activity and physiological effects more faithfully.

#### 4.1.7 Simplification of the relationship between VNS and physiological effects via eCAPs

As illustrated by Fig. 1, we suppose that pulse parameters only affect physiology via the evoked neural activity. This would suggest that modelling physiological responses is more straightforward from neural activity than from pulse parameters. This is illustrated Fig. 13. On the one hand, the left-hand pane shows the complexity of the mapping between current and stimulation location and laryngeal muscle activation. The current has a non-linear influence, which is different depending on the stimulation location. On the other hand, there is a strong linear relationship between the same physiological effect and A*β*-fibre activation, regardless of the underlying current or location of each stimulation.

### 4.2 VNS dosing for clinical applications

#### 4.2.1 VNS dosing using neural recordings

The high variability of physiological effects (Section 3.1) as well as neural activations (Section 3.3) between subjects in the response to VNS suggests that personalized dosing is needed to optimize VNS settings for a desired therapeutic effect while avoiding side effects. Furthermore, the results in 3.2 suggest that existing dosing methods that adjust a single parameter, most often current, or cycle through preset programs are unlikely to lead to optimal settings. Stimulation parameters all influence (to a greater or lesser extent) the physiological responses of the subject. While per sonalized high-dimensional parameter search will likely lead to more precise physiological responses, the curse of dimensionality makes brute force search techniques impractical in the operating room or clinic.

We observe that neural recordings and the distinction between pulse and train parameters described in 3.3 and 3.4 and illustrated Fig. 1 can help tackle this challenge. Considering the pulse parameters separately from the train parameters while recording neural responses would simplify and clarify where variability is occurring in patients’ responses to VNS: the variability of the neural responses is captured by pulse parameters, while the variability of physiological responses is captured by train parameters and the evoked neural activity. On the one hand, pulse parameters can be explored with the aim of optimizing for a preferred fibre activation profile, which is informed a priori by known organ functions of the different fibre types and can be refined during dosing based on alignments with physiological effects as demonstrated in Section 3.4. For example, if the neural fulcrum introduced by Ardell et al. (2017) is a desired physiological target while trying to mitigate breathing effects, an optimization objective would be to activate B-fibre while minimising A*γ*-fibre activation. On the other hand, train parameters provide additional levers for controlling the depth of physiological effects and trade-offs between on- and off-target effects, as illustrated in Fig. 6.

With such a large parameter space an algorithmic approach to dosing would be critical to take full advantage of neural recordings in VNS dosing. While the pulse parameter space is high-dimensional, evoked neural activity has an intrinsic timescale in the ms range and can be explored with limited physiological effects. This means neural responses can be optimized quickly and accurately across a large pulse parameter space. However, this would be highly challenging for manual approaches—as commonly used in current dosing procedures—which cannot take full advantage of the speed of neural responses and the high-dimensional pulse parameter space. Instead, an algorithmic approach to dosing would be able to consider high number of input parameters and efficiently search through multiple dimensions simultaneously. By handling the combination of neural and physiological responses to both pulse and train parameters, such an algorithm could guide the dosing procedure with optimal personalised stimulation parameter suggestions. In a companion paper^1^ we describe, implement and test such an algorithmic approach to dosing, relying on Bayesian optimization, enabling an efficient, safe, and traceable optimization of neural and physiological responses to VNS.

#### 4.2.2 Clinical relevance

Using neural recordings during VNS dosing offers the benefits of optimizing over a wide range of VNS doses in a short amount of time, improving responder rates and enabling therapies to be reoptimized over time.

Traditional dosing procedures rely on the observation of VNS-induced physiological effects to assess the effectiveness of a given set of stimulation parameters. Such physiological changes might happen in the order of a couple of seconds or minutes—e.g. in the case of short-term heart rate effects— but can take as long as hours for some autonomic function such as inflammatory responses which require blood samples to be collected. In contrast, neural activity is measured in a matter of milliseconds following a stimulation pulse, and the consistency of eCAPs across pulses of the same stimulation train as illustrated Fig. 8 suggests that evoked neural activity can be assessed with as little as a single pulse. As a result, the neural responses to hundreds of potential stimulation parameters can be explored in the same amount of time required to assess the physiological effects of a single set of stimulation parameters.

In regard to responder rates, the organisation of fascicles within the vagus nerve varies greatly across patients, and is believed to be one of the main explanations for the high rates of non-responders to current VNS therapies. Approaches to tackle this include live imaging of the nerve in surgery via Electrical Impedance Tomography (EIT) (Ravagli et al., 2020) or ultrasound (Curcean, Rusu, and Dudea, 2020) to determine the best location for the stimulation cuff. However, these approaches add to surgical complexity, and do not account for differences in the cuff-nerve interface. Instead, the fast exploration and optimization of the mapping between pulse parameters and evoked neural responses allows one to capture the neural phenotype of each patient efficiently and safely.

Finally, using neural responses during VNS dosing would allow to reoptimize VNS therapies with less stress to the patient. While not used for VNS dosing today, IPGs with neural recording capabilities or minimally invasive techniques, such as microneurography (Ottaviani et al., 2020; Verma et al., 2022), could enable neural recordings to be used periodically as efficacy decreases in response to electrode movement, scarring, or habituation.

### 4.3 Limitations

#### 4.3.1 Causal link between eCAPs and physiological effects

It is a basic assumption of VNS that eCAPs elicited by VNS lie on the causal path from stimulations to their physiological effects. However, eCAPs actually responsible for causing a specific physiological effect might elude recording. On the other hand, eCAPs recorded from a specific site on the VN might not be the ones involved in causing that physiological effect. Simple correlation, the co-occurrence of eCAPs with certain effects, is not necessarily causation.

Traditional causal inference based on statistical principles (Pearl, 2009), which allows us to establish causal connections from observed data, is not applicable either. Such inference requires deliberate or gratuitous randomization on the process level. However, as we observed in this study, the relationship between stimulation parameters and eCAP patterns is essentially deterministic without any process noise.

Related methods for establishing a link between a fibre type activation and a physiological effect are based on correlation or regression analysis (Qing et al., 2018; Chang et al., 2020). Such methods typically depend on the correct choice of a regression model. For example, relying on correlation or a linear regression model for the typically highly nonlinear relationship between fibre activations and physiological responses can produce misleading results.

Therefore we used a weaker criterion to establish a possible causal link between certain eCAPs and a physiological effect: an alignment of DRCs of eCAP activations and response. The criterion requires that points of onset and saturation of the two DRCs agree. This criterion cannot prove a causal link but only suggest plausible candidates that might deserve further investigation. While similar approaches have been applied implicitly in the literature (for example in Verma et al. (2022) and Blanz et al. (2022)), here we have explicitly stated the principle and have applied it systematically in our data analysis, for example in Section 3.4.

#### 4.3.2 Limitations of recorded evoked neural activity

In this study, we use a simple bound-based metric to measure the activation of individual neural fibres. However, this might give an incomplete view of the true evoked neural activity. Firstly, cuff electrodes only measure the neural activity in their vicinity. Secondly, fascicles with similar conduction velocities can appear as a single eCAP in the neurograms. A fine understanding of the true evoked neural activity is critical to predict and optimize downstream physiological effects. Computational modelling of the responses of neurograms from different radial locations could help uncover the geometry of the nerve and its evoked activity, and will be studied in future works.

#### 4.3.3 Limited number of subjects

Given the high-dimensionality of the entire parameter space, as well as the time constraints imposed by surgery, broader exploration was preferred to consistency between subjects. We argue that if individual stimulation parameters have a significant importance in the dosing of a single subject, then they might be worth considering during the dosing of other subjects.

#### 4.3.4 Translation from anaesthetised to awake subjects

The anesthetic agent used during VNS was propofol, which maintained a stable depth of anesthesia for recording vagal nerve activity. Throughout our experiments, anesthesia had little effect on recorded evoked neural activity. However, it certainly had an effect on the physiological response. Propofol is known to cause a fall in arterial pressure and a reduction in heart rate caused by central baroreflex sensitization (Whitwam et al., 2000). As a result, the integration of eCAPs into physiological responses is expected to vary. The relationship between B-fibres and bradycardia may be less obvious in the awake state and tachycardia induction may occur at smaller currents than the ones reported. The translation of the mechanisms presented here to the awake state will be addressed in future work.

#### 4.3.5 Translation from porcine to human models

The approach taken by this study is non-destructive and could be used on humans to refine the understanding of the mechanisms of action of VNS in human subjects. While pig and human vagus nerves are very similar in diameter and composition (Pelot et al., 2020), making the porcine model the most suitable for clinical translation, morphological differences exist. The porcine vagus contains on average 10 times more fascicles than a human’s (Pelot et al., 2020). As a result, we hypothesise that understanding evoked neural activity from neurograms and their relationship to physiological responses might be more challenging in porcine subjects than humans.

#### 4.3.6 Limitations of the stimulation parameter space

The range of pulse widths and frequencies explored in this work covered ranges used in approved VNS therapies. For example, LivaNova, 2020 uses pulse widths up to 500 µs and frequencies up to 30 Hz. Higher frequencies in the order of 100 Hz and 1kHz were not studied in this work, but have demonstrated promising properties for directionally-and fibre-selective stimulations (Patel and Butera, 2015; Patel et al., 2017; Chang et al., 2022). The current was explored up to 2.5 mA due to hardware constraints. Neural dose response curves show most neural fibres were past or close to saturation at this current (e.g. Fig. 9a). However, this maximum ‘current was likely not enough to elicit any C-fibre responses, which have been observed for much higher currents or with specialist preparations (Tosato et al., 2006; Yoo et al., 2013). The train duration explored in this study (1-10 s) were shorter than those typically used in current VNS therapies (e.g. 30 s in LivaNova, 2020). However, strong physiological effects were observed within this range of train durations without the need for longer durations that could potentially compromise the stability of the subject. We believe longer train durations as commonly used in current VNS therapies might not be necessary to achieve clinical effectiveness and might even be detrimental to the therapy by making side-effects more prevalent.

#### 4.3.7 Limitations of the optimization in the eCAP space

In this work we only consider squared stimulation pulses. As studied Section 3.3, this means that the evoked neural activity only depends on current amplitude, pulse width and stimulation location. However, this might limit the extent to which specific fibres can be activated while mitigating others: while pulse width and stimulation location can be optimized to preferentially target B-fibres, further selectivity might be required to fully mitigate other fibres with significantly lower neural thresholds such as fast A-fibres. A possible solution could be to use non-squared pulse shapes, which have demonstrated promising results towards selective fibre activation (Vuckovic, Tosato, and Struijk (2008), Pěclin and Rozman (2014), Dali et al. (2018), Dali et al. (2019)). Although more complex pulse shapes would bring additional pulse parameters to consider, the speed and safety advantages of optimizing within the eCAP space would allow to deal with this increase in dimensionality of the input space.

#### 4.3.8 Adaptability of eCAPTURS Framework

The key results of this study informed the design of the eCAPTURS framework depicted in Fig 1A. This framework can guide one’s understanding of neural and physiological mechanisms of VNS. While the version of this framework depicted in Fig. 1 is reflective of the specific experimental set-up described in Section 2, it can easily be adapted for experimental set-ups that utilize different VNS settings.

One example of how the framework could be adapted is by incorporating new pulse parameters from complex wave forms such as a depolarizing pre-pulse as studied by (Vuckovic, Tosato, and Struijk, 2008) or chopped pulses as studied by Qing, Ward, and Irazoqui (2015) and Dali et al. (2019). The latter case is a particularly interesting use of the framework as the intra-burst frequency parameter used in chopped pulses would be considered a pulse parameter as it affects evoked neural activity by leveraging the refractory periods of different fiber types, while interburst frequency would remain a train parameter as it does not significantly affect evoked neural activity.

Another example of how the framework could be adapted is the inclusion of a closed-loop control mechanism (Bilgutay et al., 1968; Tosato et al., 2006; Ugalde et al., 2014; Ojeda et al., 2016; Mylavarapu et al., 2023). These studies use live monitoring of the RR interval as an input for determining the timing and/or parameters for stimulating the nerve. In this case, specific delays from R peaks in the ECG as seen in studies by Ojeda et al. (2016) and Ugalde et al. (2014), could be included in the framework as ”train parameters” as timing of the stimulation impacts physiological effects but is not expected to influence evoked neural activity. In this example, an arrow would likely connect the ”Physiological Context” box and the ”Train Parameter” box depicted in Fig 1A.

In addition to adding new stimulation parameters, studying VNS for other chronic conditions would require adding or removing physiological effects. The physiological effects listed in Fig 1 are those that are most relevant to VNS for HF. Other physiological effects of VNS that might be commonly considered include dysphonia (Ardesch et al., 2010), biomarkers of inflammation (Kwan et al., 2016), and muscle activity in the gut (Lu et al., 2018). As we continue to deepen and broaden our experimental work, this framework will be extended to capture a more complete picture of the relationship between VNS, neural effects, and physiological changes.

## 5 Conclusion

Choosing the right stimulation parameters in VNS has been one of the primary obstacles impeding its wider application to treat various chronic diseases. In this study, we investigate the relationship between VNS parameters, neural responses and short-term physiological responses to stimulation through exploring a wide range of systematic and extensive sets of VNS parameters. We find that there is high inter-patient variability which necessitates personalized VNS dosing, and that increasing the range of stimulation parameters considered in VNS dosing improves outcomes; however, this comes with the cost of more extensive and time-consuming procedures, which are impractical under clinical constraints. We propose that evoked neural activity can be used to mitigate this shortcoming as it can be explored much faster and more safely than physiological effects by focusing on pulse parameters, while being closely associated to downstream physiological effects via train parameters. Dosing strategies harnessing insights from evoked neural activity could explore and optimize over the stimulation parameter space more thoroughly and in a shorter amount of time as existing dosing procedures, leading to better therapeutic outcomes in fewer clinic visits for the patient. Together, these findings suggest that novel calibration methods leveraging recordings of evoked neural activity may enable a new kind of precision medicine with personalized, safe and efficient VNS dosing. Future works will attempt to demonstrate and validate such calibration procedure in a pre-clinical setting.

## Authors contribution

AB and LW analyzed the data with contributions by OT-L. AB, LW, MS prepared the first draft of the manuscript. MT made substantial further contributions to the manuscript. PF-P, OA, MS, TE designed the animal experiments. PF-P and JM guided the animal experiments. BP, MJ designed and implemented the cloud based data processing and storage pipeline. BA, SG, MJ designed and implemented the NeuroTool interface. SS, SG, PG, MP, MS, BA, MJ designed and built the BIOS Neural Interface. LA, ES, SL, GL contributed to reviewing and rewriting of the manuscript. EH, AT, JJ, OA, TE conceived of the study. OA supervised the study.

## Footnotes

1 Currently under review.

